# Btbd11 is an inhibitory interneuron specific synaptic scaffolding protein that supports excitatory synapse structure and function

**DOI:** 10.1101/2021.11.01.466782

**Authors:** Alexei M Bygrave, Ayesha Sengupta, Ella P Jackert, Mehroz Ahmed, Beloved Adenuga, Erik Nelson, Hana L Goldschmidt, Richard C Johnson, Haining Zhong, Felix L Yeh, Morgan Sheng, Richard L Huganir

## Abstract

Synapses in the brain exhibit cell-type-specific differences in basal synaptic transmission and plasticity. Here, we evaluated cell-type-specific differences in the composition of glutamatergic synapses, identifying Btbd11, as an inhibitory interneuron-specific synapse-enriched protein. Btbd11 is highly conserved across species and binds to core postsynaptic proteins including Psd-95. Intriguingly, we show that Btbd11 can undergo liquid-liquid phase separation when expressed with Psd-95, supporting the idea that the glutamatergic post synaptic density in synapses in inhibitory and excitatory neurons exist in a phase separated state. Knockout of Btbd11 from inhibitory interneurons decreased glutamatergic signaling onto parvalbumin-positive interneurons. Further, both *in vitro* and *in vivo*, we find that Btbd11 knockout disrupts network activity. At the behavioral level, Btbd11 knockout from interneurons sensitizes mice to pharmacologically induced hyperactivity following NMDA receptor antagonist challenge. Our findings identify a cell-type-specific protein that supports glutamatergic synapse function in inhibitory interneurons—with implication for circuit function and animal behavior.

## INTRODUCTION

The postsynaptic density (PSD) of glutamatergic synapses in the brain is a densely packed protein rich structure that supports excitatory synaptic transmission and synaptic plasticity. Presently, our understanding of the PSD largely comes from studying excitatory neurons (ENs) which make up 80-90% of neurons in cortical and hippocampal circuits (Hu et al., 2014). Impaired function of inhibitory interneurons (INs) is associated with psychiatric disease and neurological disorders (Lewis et al., 2005; Lisman et al., 2008; Marín, 2012). More specifically, impaired glutamatergic excitation of INs, particularly parvalbumin (PV) positive INs, is linked to the pathophysiology of schizophrenia (Lewis et al., 2005; Lisman et al., 2008). PV-INs have been extensively studied, and their role in regulating circuit activity (*i*.*e*., controlling the precise timing of EN cell firing), supporting rhythmic neuronal activity (*i*.*e*., generation of gamma oscillations), and controlling animal behavior is well documented (Cardin et al., 2009; Donato et al., 2013; Kuhlman et al., 2013; Mann et al., 2005; Sohal et al., 2009). Interestingly, both basal glutamatergic transmission and plasticity in INs (including PV-INs) is different to in ENs (Chang et al., 2010; Geiger et al., 1995; Lamsa et al., 2007; Matta et al., 2013). However, very little is known about the molecular composition of the IN glutamatergic post-synaptic density (inPSD) and whether cell-type-specific specializations exist to govern these distinct properties.

Recent evidence indicates that many components of the glutamatergic PSD undergo liquid-liquid phase separation (LLPS), suggesting the PSD likely exists as a phase-separated structure that aids synapse function (Chen et al., 2020; Feng et al., 2019; Zeng et al., 2016, 2019). LLPS could be particularly important for the stability and function of inPSD’s, which are frequently located directly on the dendritic shaft of INs (Hu et al., 2014) rather than being nested within dendritic spines as is common in ENs (but note Keck et al., 2011 and Sancho et al., 2018). Molecular mechanisms must support PSD stability, as shaft glutamatergic synapses located within PV-INs are on average more stable than their counterparts in INs when assessed with longitudinal imaging *in vivo* (Melander et al., 2021). Here, we use a combination of genetic tools and proteomics to identify cell-type-specific PSD proteins. We characterize Btbd11 as a novel inPSD which undergoes LLPS and regulates glutamatergic synapses in INs as well as neuronal circuit function.

## RESULTS

### Identification of Btbd11 as a novel inPSD protein

To gain insights into the molecular composition of inPSDs, we bred conditional Psd-95-GFP knockin mice (second generation mice based on Fortin et al., 2014) with vGAT^Cre^ or CaMKII^Cre^ animals, tagging a core scaffold protein at glutamatergic synapses in INs or ENs, respectively. We collected cortex and hippocampal tissue from these animals and performed GFP-immunoisolation experiments (**Figure 1A**). To compensate for the lower abundance of INs we pooled samples from 4 vGAT^Cre^:Psd-95-GFP mice to compare with 1 CaMKII^Cre^:Psd-95-GFP mouse. Utilizing mass spectrometry and label free quantification, we identified immuno-isolated PSD complexes preferentially originating from INs or ENs. Normalising to the amount of Psd-95 we found proteins that were specifically enriched in INs (**Figure 1B, Table S1**). Such proteins were considered putative inPSD proteins. The top hit was Btbd11, an ankyrin repeat and BTB/POZ domain-containing protein.

**Figure 1.**
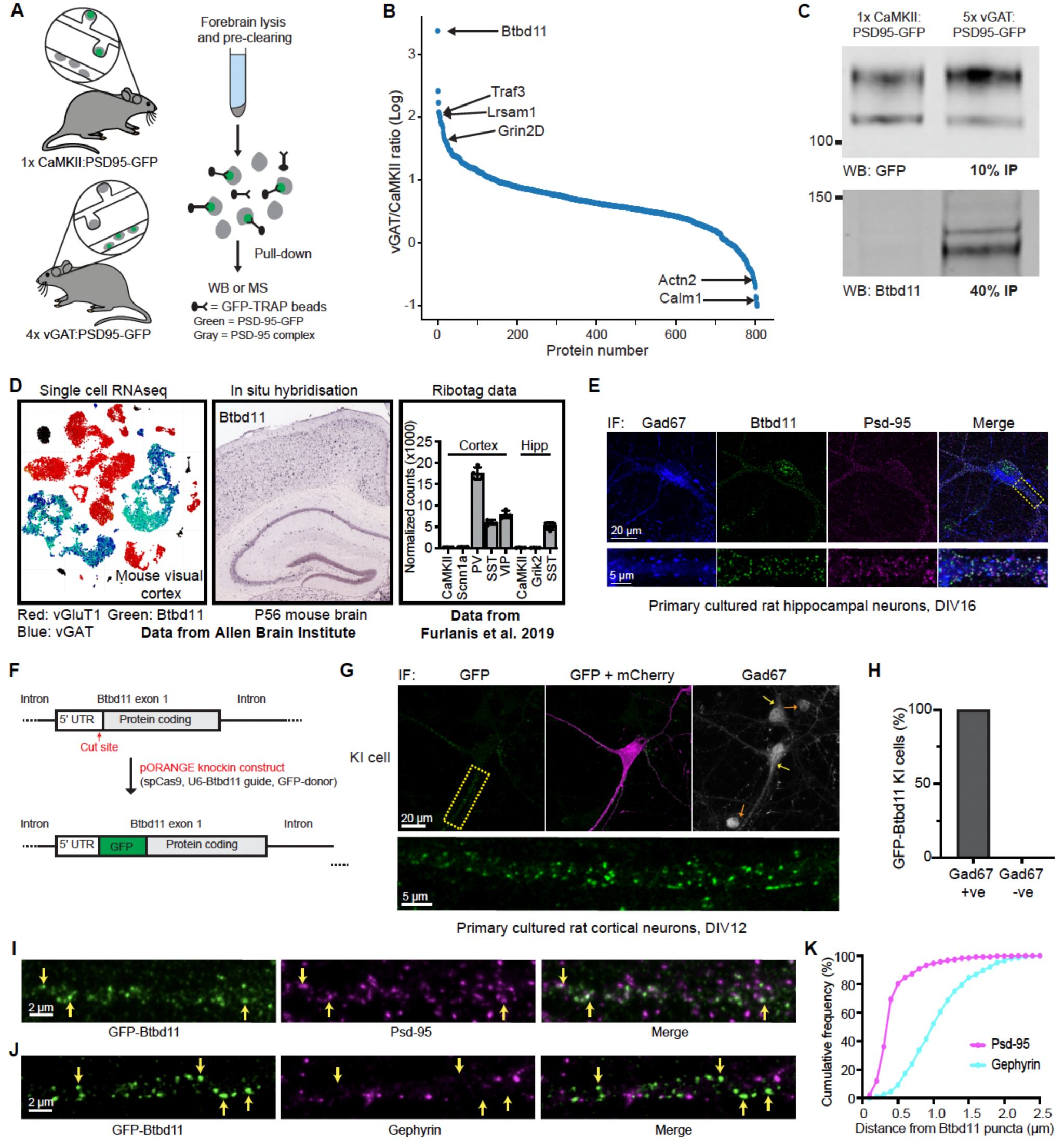
Identification of Btbd11 as a novel inPSD protein. (A) Schematic of Psd-95-GFP knockin in CaMKII or vGAT positive neurons to label excitatory and inhibitory neurons, respectively, for use in immunoisolation experiments. (B) Proteins identified using semi-quantitative mass spectrometry following PSD-95-GFP immunoisolation expressed as a vGAT/CaMKII ratio. Before calculating the ratio, proteins were normalized to levels of Psd-95. (C) Psd-95-GFP immunoisolation with CaMKII:Psd-95-GFP or 5x vGAT:Psd-95-GFP (pooled) mice followed by western blots for GFP (detecting Psd-95-GFP, the “bait”) from 10% of the pull-down, or Btbd11 (a candidate inPSD) from 40% of the pull-down. (D) Assessment of different RNA databases showing that Btbd11 RNA is highly enriched in INs in the cortex and hippocampus. (E) Immunofluorescence in rat primary hippocampal neurons (DIV16) for Gad-67, Btbd11 and Psd-95. The region in the yellow dashed box is enlarged below. (F) Schematic of the CRISPR knockin strategy used to label endogenous Btbd11. GFP was targeted to the N-terminal region of Btbd11. (G) Example of GFP-Btbd11 knockin cell from primary cultured rat cortical neurons electroporated with knockin constructs and a mCherry cell-fill. Immunofluorescence was used to boost the GFP signal (green) and to identify Gad-67 positive cells (a marker of INs). Yellow arrows indicate Gad-67 positive cells (with observed cytoplasmic signal) and orange arrows indicate Gad-67 negative cells (the was some non-specific nuclear signal). (H) All of the 31 GFP-Btbd11 knockin cells were also Gad-67 positive, showing the specificity of Btbd11 in INs. (I and J) Zoom in along the dendrites of GFP-Btbd11 knockin cells with immunofluorescence for Psd-95 or Gephyrin, respectively. Yellow arrows indicate a subset of Btbd11 puncta. (K) Cumulative frequency plots indicating the puncta-to-puncta distance of Btbd11 with Psd-95 (magenta) or Gephyrin (blue).

Btbd11 was previously identified in both Psd-95 pull-down and proximity labelling experiments (Fernández et al., 2009; Uezu et al., 2016), but its cell-type-specific expression pattern was not explored. Furthermore, a biological function of Btbd11 remains completely unknown, in the brain or otherwise. We generated and validated an antibody against Btbd11 (**Figure S1**) as no commercial antibodies were available. To confirm our mass spectrometry results, we repeated GFP-immuno-isolation experiments and used western blots to confirm that Btbd11 was selectively pulled-down from IN samples (**Figure 1C**). We then explored published RNA datasets and found that Btbd11 mRNA expression is exclusive to INs in the cortex and hippocampus of mice (Furlanis et al., 2019; Tasic et al., 2016) (**Figure 1D**).

Using immunofluorescence and our antibody for Btbd11 in cultured rat hippocampal neurons we observed punctate Btbd11 expression which co-localized with Psd-95 in Gad67-positive INs (**Figure 1E**). As a secondary validation, and to circumvent any possible issues with non-specific antibody binding, we designed CRISPR knockin constructs to label endogenous Btbd11 using the ORANGE method (Willems et al., 2020). We opted to tag Btbd11 at the N-terminus with GFP, and electroporated rat primary cultures cortical neurons with the knockin construct alongside a mCherry cell-fill (**Figure 1F**). We observed a sparse population of cells with punctate GFP signal which, as expected, also were positive for Gad67 (**Figure 1G**). Indeed, of the 31 KI cells imaged 100% were also Gad67-positive (**Figure 1H**), confirming the IN-specific expression of Btbd11 in cortical neurons. We next coupled this CRISPR knockin approach with immunofluorescence to evaluate if Btbd11 puncta were enriched at synapses. GFP-Btbd11 puncta overlapped with Psd-95 puncta, indicating the protein is found at glutamatergic synapses (**Figure 1I**). In contrast, there was very little overlap with Gephyrin, a marker of GABAergic inhibitory synapses (**Figure 1J**). We quantified the relative enrichment at glutamatergic synapses by calculating the distances between GFP-Btbd11 puncta and Psd-95 or Gephyrin puncta and found that the puncta-to-puncta distance was considerably smaller at glutamatergic synapses (**Figure 1K**).

Collectively, these data highlight that there are cell-type-specific differences in the composition of glutamatergic synapses and establish Btbd11 as a novel inPSD protein that is selectively located at glutamatergic synapses.

### Btbd11 contains a PDZ binding motif which interacts with PDZ1,2 of Psd-95

Btbd11 is annotated to contain 5 ankyrin repeats, a BTB domain, and region of predicted disorder (www.uniprot.org). In addition, the C-terminal amino acids are consistent with a PDZ binding motif (PBM; **Figure 2A**). Utilizing a protein structure prediction algorithms from AlphaFold (Senior et al., 2020) we observed that Btbd11 contains additional regions of potential disorder (**Figure 2B, Video S1**). Interestingly, the C-terminal region of Btbd11 is highly conserved between species, highlighting the potential importance of the PBM (**Figure 2C**). PBM’s support interactions with PDZ domain containing proteins, including the membrane-associated guanylate kinases (MAGUK) family of proteins such as Psd-95 used as “bait” in our immune-isolation experiment. We hypothesized that the PBM of Btbd11 might support a direct interaction with the PDZ domains of Psd-95.

**Figure 2.**
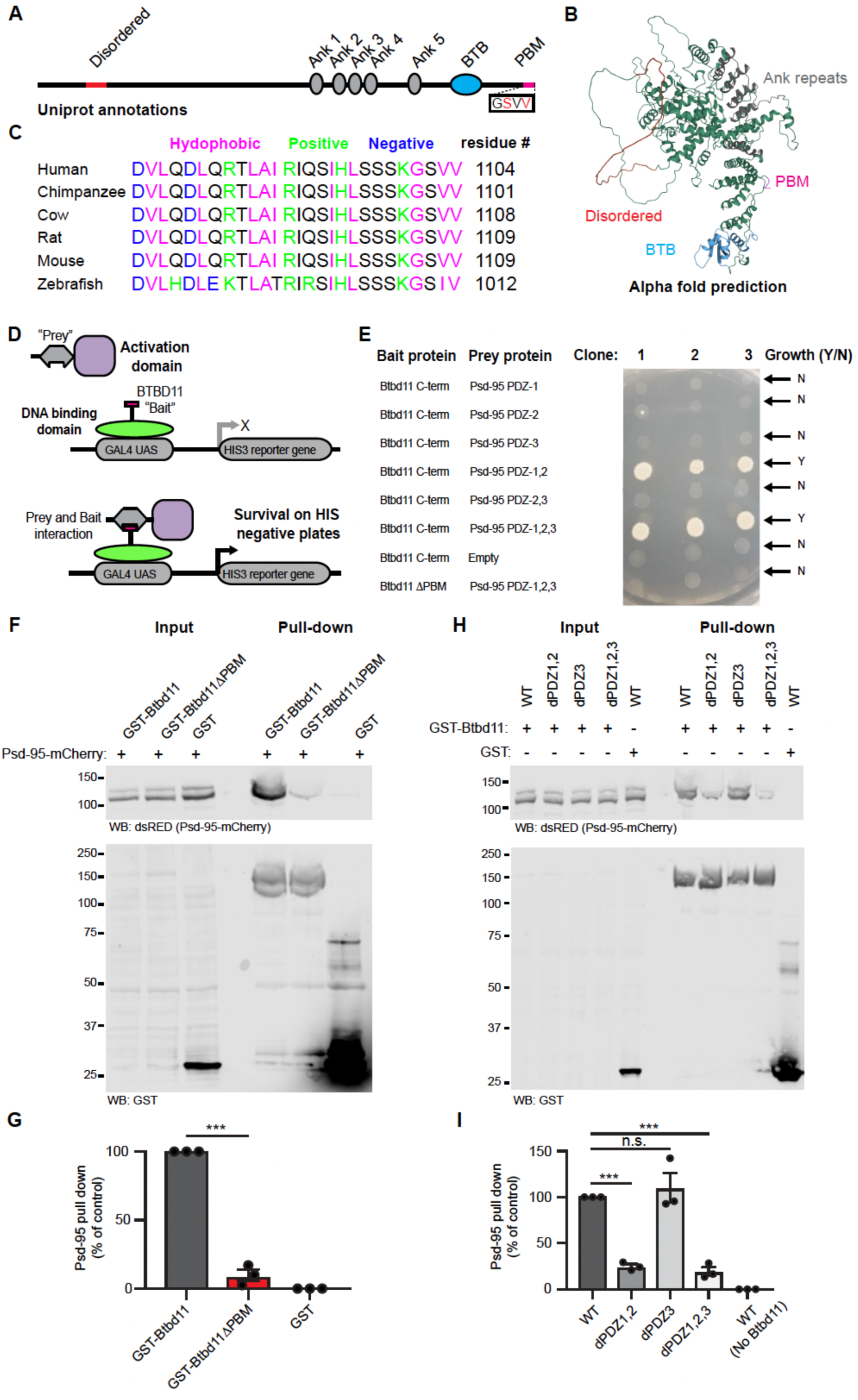
Btbd11 contains a PDZ binding motif which interacts with PDZ1,2 of Psd-95. (A) Schematic depiction of Btbd11 with annotations from Uniprot. Red = disordered, gray = ankyrin repeats (Ank), blue = BTB domain (BTB), magenta = PDZ binding motif (PDM). (B) Predicted structure of Btbd11 from AlphaFold with domains shaded as in (A). (C) C-terminal region of Btbd11 in different species showing the conservation of the PBM. (D) Schematic of targeted yeast two-hybrid experiment to assess binding of Btbd11 with different PDZ domains of Psd-95. (E) Results from targeted yeast two-hybrid experiment with growth indicating an interaction between Btbd11 and Psd-95. (F) GST pull-down experiments evaluating the ability of GST-Btbd11 or a mutant lacking the PBM (GST-Btbd11ΔPBM) to interact with Psd-95-mCherry in HEK cells. GST only was included as a negative control. (G) Qualification of Psd-95-mCherry pulled-down by GST-Btbd11 or GST-Btbd11ΔPBM. (H) GST pull-down experiments in HEK cells evaluating the ability of GST-Btbd11 to interact with Psd-95-mCherry point mutants designed to disrupt PDZ domain binding. (I) Quantification of Psd-95-mCherry mutants pulled-down by GST-Btbd11. Error bars display S.E.M. *** indicates p < 0.001. *See* **Table S2** for full statistical used for this and all subsequent analyses.

To test if there is a direct interaction between Btbd11 and Psd-95 through PBM-PDZ domain interactions we used a yeast 2-hybrid approach. We created a fusion protein consisting of a DNA binding domain and the C-terminus of Btbd11 +/- the PBM as “bait”, and used an activation domain-Psd-95 fusion protein containing different combinations of the 3 PDZ domains as “prey” (**Figure 2D**). We only observed yeast growth, a readout of a protein-protein interaction, when PDZ domains 1,2 or 1,2,3 were present (**Figure 2E**). Btbd11 lacking the PBM did not interact with PDZ 1,2,3 of Psd-95, highlighting the necessity of the PBM for binding.

To complement the yeast 2-hybrid approach we adopted a GST pull-down strategy in mammalian cells. We cloned full-length GST-Btbd11 +/- the PBM and expressed these GST fusion proteins (or GST only) in HEK cells alongside Psd-95-mCherry, then lysed the cells and performed a GST pull-down. We found that Btbd11 was able to pull-down Psd-95-mCherry, but only when the PBM was present (**Figure 2F**,**G**). To confirm the binding was mediated by PDZ1,2 of Psd-95, we cloned mutants of Psd-95-mCherry in which the individual PDZ domain function was disrupted (dPDZ1-3) based on previously generated mutants (Imamura et al., 2002). Confirming our observations in yeast, we found diminished binding to Btbd11 with Psd-95 harbored mutations in PDZ1,2 or PDZ1,2,3. Mutations to PDZ3 alone had no discernable effect on Btbd11 binding, highlighting the specificity of the interaction.

Having established a direct interaction with Psd-95, we sought to identify other potential interaction partners. We conducted a yeast 2-hybrid screen using either the BTB domain or the C-terminal region containing the PBM of Btbd11 as “bait”. From the screen with the C-terminus of Btbd11 we identified well known PDZ-containing synaptic proteins including Psd-93, Sap-102 and Pick1 as putative interactors (**Figure S2**). Using the BTB domain we pulled out Ataxin1 and Ataxin-1like as potential interactors (**Figure S2**). Although not yet tested we expect that the tandem ankyrin repeats of Btbd11 facilitate other protein-protein interactions, as is common for proteins with ankyrin repeats (Mosavi et al., 2004).

Taken together, these data show that Btbd11 contains a highly conserved C-terminal PBM which mediates a direct interaction with PDZ1,2 of Psd-95. Furthermore, our yeast 2-hybrid screen indicates that Btbd11 can interact with other key synaptic proteins, including other MAGUK family members and proteins known to regulate the trafficking of glutamate receptors (*i*.*e*., Pick1).

### Liquid-liquid phase separation of Btbd11 when expressed with Psd-95

We next sought to explore the properties of exogenously expressed Btbd11. To do this we cloned GFP and mCherry tagged Btbd11 with or without the PBM necessary for an interaction with Psd-95. In HEK cells expression of GFP-Btbd11 led to the formation of striking fibril-like structures (**Figure 3A**). Remarkably, when Psd-95-mCherry was expressed alongside GFP-Btbd11 the fibril structures were replaced with large spherical intracellular droplets with a high degree of Btbd11 and Psd-95 colocalization (**Figure 3B**,**C**). Droplet formation was critically dependent upon an interaction between Btbd11 and Psd-95, since deletion of the PBM of Btbd11 abolished their formation (**Figure 3D**,**E**). By contrast, formation of fibrils was not dependent upon the PBM (see **Figure S3** for quantification of fibril and droplet formation under different conditions).

**Figure 3.**
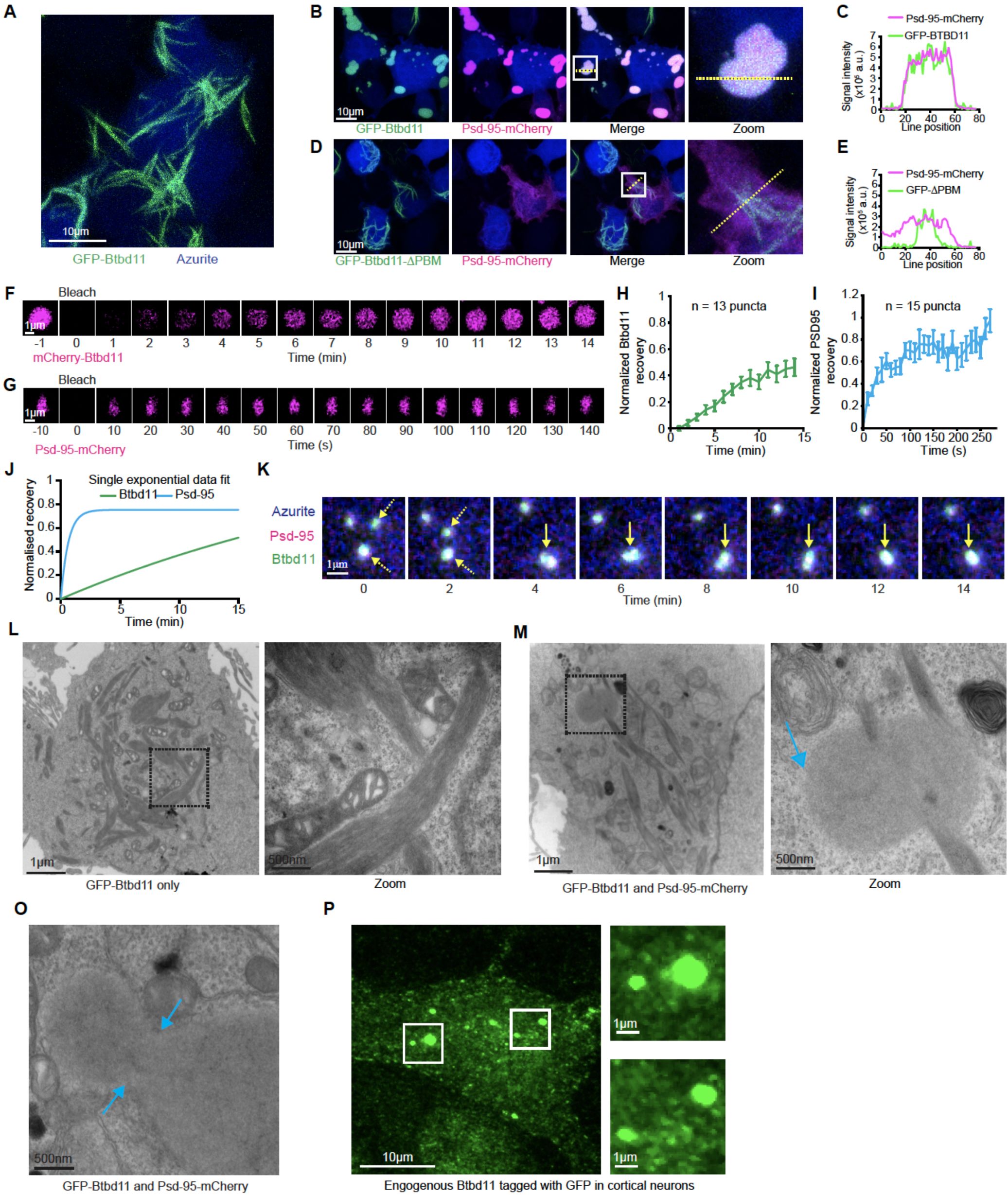
Liquid-liquid phase separation of Btbd11 when expressed with Psd-95. (A) Expression of GFP-Btbd11 (green) and an azurite cell-fill (blue) in HEK cells led to the formation of large fibril-like assemblies. Scale bar = 10μm. (B) Co-expression of GFP-Btbd11 (green) with Psd-95-mCherry (magenta) and an azurite cell-fill (blue) led to the formation of spherical droplets in HEK cells. The region within the white box is enlarged on the right (Zoom), and the yellow dotted line indicates where a line scan (C) is shown for Psd-95-mCherry and GFP-Btbd11. (D) Co-expression of GFP-Btbd11ΔPBM (green) with Psd-95-mCherry (magenta) and an azurite cell-fill (blue) in HEK cells led to the formation of fibril-like assemblies with the absence of spherical droplets. The region within the white box is enlarged on the right (Zoom), and the yellow dotted line indicates where a line scan (E) is shown for Psd-95-mCherry and GFP-Btbd11ΔPBM. (F and G) Fluorescence recovery after photobleaching (FRAP) of mCherry-Btbd11 and Psd-95-mCherry co-expressed with Psd-95-GFP and GFP-Btbd11, respectively. Note the difference in timescales for Btbd11 and Psd-95. Scale bar = 1μm. (H and I) Quantification of fluorescence recovery for Btbd11 (n = 13) and Psd-95 (n = 15), respectively. Error bars indicate S.E.M. (J) Plot of the exponential fit of FRAP data for Btbd11 (green) and Psd-95 (blue). (K) Longitudinal confocal imaging of a HEK cell transfected with GFP-Btbd11 (green), Psd-95-mCherry (magenta) and an azurite cell-fill (blue). Yellow dotted arrows indicate two puncta that come together and form a single droplet (solid yellow arrow). Scale bar = 1μm. (L and M) Electron microscope images of HEK cells transfected with GFP-Btbd11 or GFP-Btbd11 and Psd-95-mCherry, respectively. The black dotted box is enlarged on the right of each panel. Respective scale bars are indicted at the bottom left of each image. In (M) the red arrow indicates a putative droplet observed with Btbd11 and Psd-95 co-expression but not with Btbd11 expression alone. (O) An example of a putative Btbd11 and Psd-95 droplet that looks to be in the process of fusion or fission with cyan arrows indicating the neck. (P) Example of a GFP-Btbd11 rat primary cortical neuron with droplet-like assemblies in the cell body as well as puncta along the dendrites (not-shown). White boxes are enlarged on the right-hand size. Respective scale bars are indicted at the bottom left of each image.

The spherical puncta formed by Btbd11 and Psd-95 were reminiscent of biomolecular condensates arising from liquid-liquid phase separation (LLPS) of abundant PSD proteins including SynGAP and Psd-95 (Zeng et al., 2016). A hallmark of LLPS is fluorescent recovery after photobleaching (FRAP), indicating a dynamic exchange of biomolecules between putative condensates and the surrounding cytosol—a phenomena that is not expected if puncta are formed from protein aggregates locked into a solid state. To test this, we transiently expressed mCherry-Btbd11 with Psd-95-GFP (**Figure 3F**) or Psd-95-mCherry with GFP-Btbd11 (**Figure 3G**) and photobleached individual puncta in HEK cells. We observed signal recovery for both Btbd11 (**Figure 3H**) and Psd-95 (**Figure 3I**), with Psd-95 recovering at a faster rate (**Figure 3J**).

LLPS can be facilitated by proteins which contain regions of intrinsic disorder (Uversky, 2017). Btbd11 contains predicted regions of disorder within its N-terminal region (**Figure 2A**,**B**) which we speculated could promote LLPS. To test this, we cloned a GFP- and mCherry-Btbd11 that lacked a large portion of the N-terminus upstream of the ankyrin repeats (5xANK-BTB). When expressed with Psd-95, large aggregates formed in which there was also a high degree of co-localization between 5xANK-BTB and Psd-95 (**Figure S3**). However, in addition to lacking the spherical droplets observed with full-length Btbd11, the 5xANK-BTB assemblies did not show FRAP, indicating the structures were unlikely to be liquid-like assemblies (**Figure S3**).

A further hallmark of LLPS is the coalescence of puncta over time to form larger droplets. Through time-lapse imaging of HEK cells expressing GFP-Btbd11 and Psd-95-mCherry we observed fusion of puncta over time (**Figure 3K, Video S2**). Frequently purified proteins—or portions of proteins—are mixed *in vitro* to validate LLPS. Our attempts to purify Btbd11 to conduct such experiments have so far been unsuccessful, likely because Btbd11 readily forms fibrils which make protein purification challenging. Therefore, to get an orthogonal validation that the puncta were true “membrane-less organelles” we performed electron microscopy on HEK cells expressing either GFP-Btbd11 alone, or in combination with Psd-95-mCherry. As expected, considering on our fluorescent imaging, in the Btbd11-only condition we found cells with fibril-like structures (**Figure 3L**). We also observed fibril-like structures when Btbd11 was expressed with Psd-95, but also identified droplet-like assemblies in the cytoplasm, which occasionally also appeared to be impaled by the fibrils (**Figure 3M**). Critically, these droplets appeared electron dense but clearly lacked any kind of lipid bilayer. Furthermore, we found examples of droplets that looked like they could be in the process of fusion (**Figure 3O**).

While these data came from exogenously expressed Btbd11 in HEK cells we did observe putative droplets in a CRISPR knockin cell in which endogenous Btbd11 was tagged with GFP in primary cortical neurons (**Figure 3P**). Thus, under certain conditions, endogenous Btbd11 might form biomolecular condensates at the cell body, in addition to being enriched at the synapse (**Figure 1G**).

These data strongly indicate that Btbd11 can undergo LLPS with Psd-95 in living cells, identifying for the first time an inPSD-specific protein which promotes phase separation. Since LLPS at the PSD is suggested to play important roles in synapse function, the ability of Btbd11 to promote LLPS of Psd-95 (expressed alone Psd-95 does not form intracellular droplets) could have important consequences on synapse function.

### Exogenous expression of Btbd11 stabilizes Psd-95 at glutamatergic synapses

Having observed the striking properties of exogenously expressed Btbd11 in HEK cells, we wondered about the consequence of Btbd11 overexpression in neurons. To assess this, we expressed GFP-Btbd11 in rat hippocampal neurons. Large fibril-like structures formed in the cell body and dendrites of transfected cells (**Figure 4A**). These fibril structures were stable, as observed by a lack of FRAP when a portion of the fibril was bleached (**Video S3**). GFP-Btbd11 was also observed at glutamatergic synapses when imaged with higher magnification, as indicated by co-localization with Psd-95, but not gephyrin (**Figure 4B**), confirming our previous observations with endogenously tagged Btbd11 (**Figure 1I-K**). We speculated that Btbd11’s PBM would be necessary for proper synaptic targeting and confirmed this by showing there was no synapse localization of Btbd11 that lacked a PBM (**Figure 4C**,**D**).

**Figure 4.**
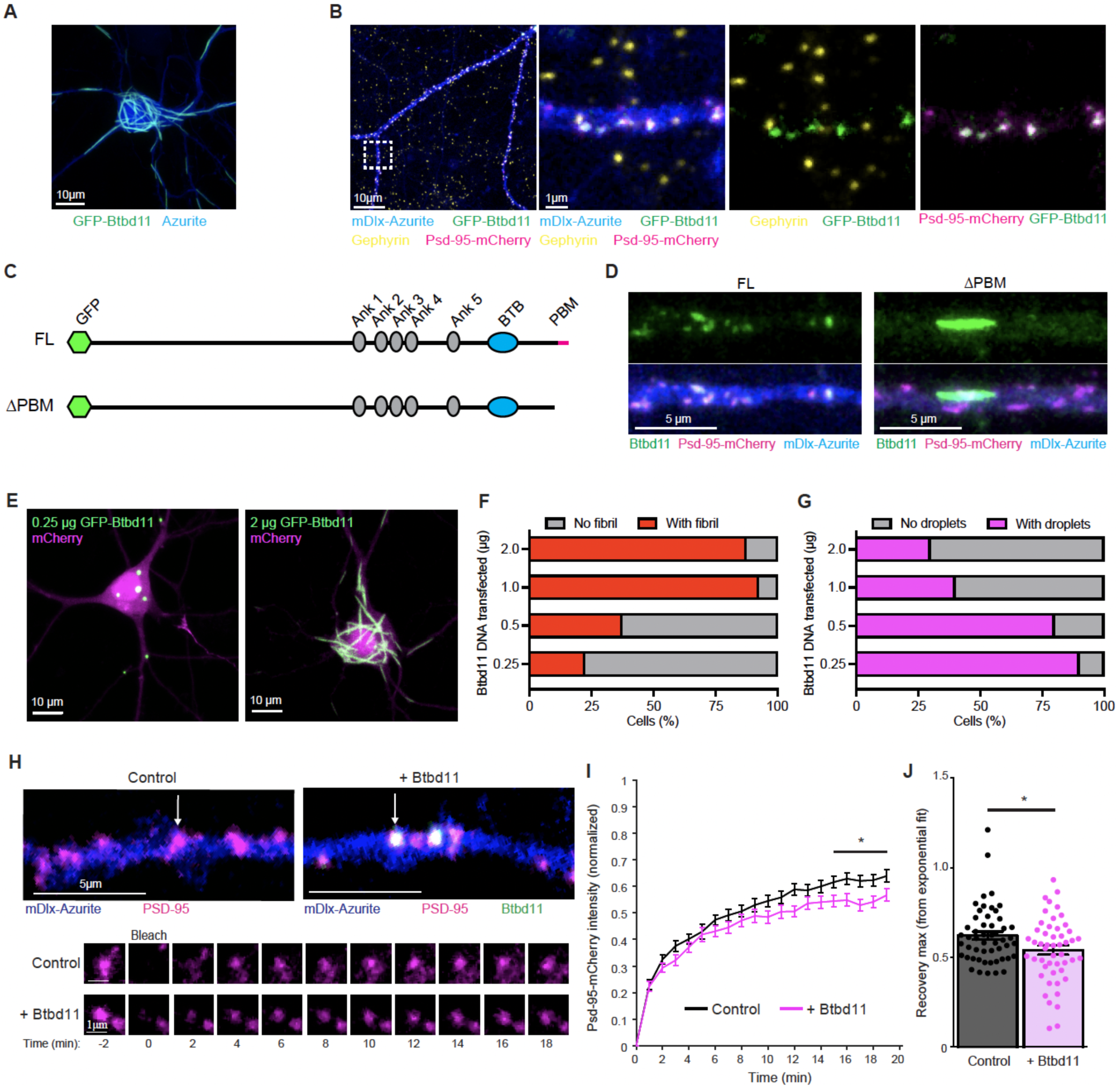
Exogenous expression of Btbd11 stabilizes Psd-95 at glutamatergic synapses. (A) Confocal image from a primary cultured rat hippocampal neuron transfected with GFP-Btbd11 (green) and azurite as a cell-fill (blue) with prominent formation of fibril-like assemblies. Scale bar = 10μm. (B) Confocal image from a primary cultured putative IN transfected with GFP-Btbd11 (green), Psd-95-mCherry (magenta), mDlx-Azurite (blue) with immunohistochemistry for Gephyrin (yellow). The region within the white dotted box is enlarged on the right with different combinations of channels. Scale bars in the bottom left of the images. Note the lack of co-localization between GFP-Btbd11 and Gephyrin. (C) Schematic of GFP-Btbd11 and GFP-Btbd11ΔPBM constructs. (D) Confocal image of putative primary cultured hippocampal interneurons, identified with an azurite cell-fill (blue) under the control of the mDlx enhancer to drive expression in INs. Full-length GFP-Btbd11 (FL, left) and GFP-Btbd11ΔPBM (ΔPBM, right) shown in green was co-expressed with Psd-95-mCherry (magenta). Synaptic puncta were observed with the FL but not ΔPBM Btbd11 construct. Scale bar = 5μm. (E) Confocal image from transfected primary cultured rat hippocampal neurons with varying amounts of GFP-Btbd11 (green) and mCherry as a cell-fill (magenta). Scale bar = 10μm. (F and G) Quantification of the proportion of cells with fibrils (red bars) or droplets (magenta bars), respectively, when GFP-Btbd11 is transfected in different quantities. (H) Live-cell confocal imaging and FRAP experiments in putative INs (identified with mDlx-azurite) in which Psd-95-mCherry (magenta) is bleached when expressed alone, or in the presence of Gfp-Btbd11 (green). The lower panels show FRAP of the individual puncta labeled in the upper panels with a white arrow. Respective scale bars are indicted at the bottom left of each image. (I) Quantification of Psd-95-mCherry FRAP under control conditions (black) or with overexpression of Btbd11 (magenta). Error bars display S.E.M. (J) Quantification of the estimated recovery maximum from an exponential fit of the FRAP data for each bleached punctum. Control data shown in black and Btbd11 overexpression data in magenta. Error bars display S.E.M. * indicates p < 0.05.

The formation of fibril-like structures is common to proteins which also undergo LLPS, with liquid to solid phase transitions occurring at saturating concentrations (Alberti and Dormann, 2019; Molliex et al., 2015). We tested if fibril formation was dose-dependent by expressing increasing amounts of GFP-Btbd11 DNA (0.25 – 2μg/well) alongside a mCherry cell fill (**Figure 4E**). While there was variability in the amount of DNA transfected within each transfection dose, clear patterns emerged whereby at high doses almost all mCherry positive cells contained Btbd11 fibrils (**Figure 4F**). Interestingly, at the lower doses we observed many cells that contained small droplet-like assemblies reminiscent of the LLPS condensates observed in HEK cells (**Figure 4G**; and see **Figure 3**). We speculate that these droplets form when Btbd11 interacts with endogenous Psd-95, and that the fibrils form when a concentration threshold is surpassed. As fibril structures were not observed with our endogenous CRISPR labeling of Btbd11 (at least under the basal conditions evaluated) we imagine that fibril formation is unlikely to be physiologically relevant to cellular function—but highlights intriguing properties of the protein.

The stability of Psd-95 is the same in INs as in ENs, despite INs generally lacking dendritic spines (Fortin et al., 2014). Indeed, we confirmed these observations with FRAP of overexpressed Psd-95-mCherry in putative INs or ENs (**Figure S4**). It has been postulated that biomolecular condensation through LLPS at the synapse might promote the stability of densely packed PSD proteins against the forces of Brownian motion (Chen et al., 2020; Feng et al., 2019). We suspect that inPSDs have protein specializations to promote their stability and function. Could Btbd11 promote the stability of Psd-95 at the synapse, possibly by driving LLPS? To explore this possibility, we expressed Psd-95-mCherry with or without GFP-Btbd11 in putative INs in hippocampal cultures identified with an Azurite cell-fill under control of the mDlx enhancer (Dimidschstein et al., 2016). Psd-95 stability was monitored by FRAP of Psd-95-mCherry puncta and the recovery curves compared with or without addition of GFP-Btbd11 (**Figure 4H**). Psd-95-mCherry FRAP was slower in the presence of overexpressed Btbd11, indicating a larger immobile fraction—or a stabilizing effect on Psd-95 at the synapse (**Figure 4I**,**J**). Intriguingly, when Btbd11 was overexpressed in ENs (which normally totally lack Btbd11), no stabilization of Btbd11 was observed (**Figure S4**). These data could be explained by the fact that ENs express high levels of known Psd-95 interaction proteins (such as SynGAP) rendering additional Btbd11-dependent stabilization redundant. Together, these data show that exogenously expressed Btbd11 forms intracellular droplets and fibrils in a dose-dependent manner, and that Btbd11 can stabilize Psd-95 at the synapse in a cell-type specific manner. It is possible that the stabilizing effect on Psd-95 results from Btbd11 promoting LLPS at the PSD.

### Btbd11 KO reduces glutamatergic signaling in PV-INs

We next explored the effects of Btbd11 loss-of-function though genetic knockout by generating Btbd11 gene trap mice using IVF and frozen sperm from the European Mutant Mouse Archive (www.infrafrontier.eu). The gene trap mice did not show reduced levels of Btbd11 mRNA or protein (**Figure S5**), probably due to a truncation previously reported in the gene-trap cassette of this line (Ryder et al., 2013). By crossing the gene-trap mice with a constitutive Flp recombinase line we converted the gene-trap mice into conditional knockout animals (Btbd11^F/F^). We validated Btbd11 knockout using primary cultured cortical neurons from Btbd11^F/F^ mice with AAVs delivering GFP or GFP-Cre. There was reliable loss of Btbd11 mRNA and protein in the presence of GFP-Cre, observed with Northern blots and western blots, respectively (**Figure 5A**,**B**).

**Figure 5.**
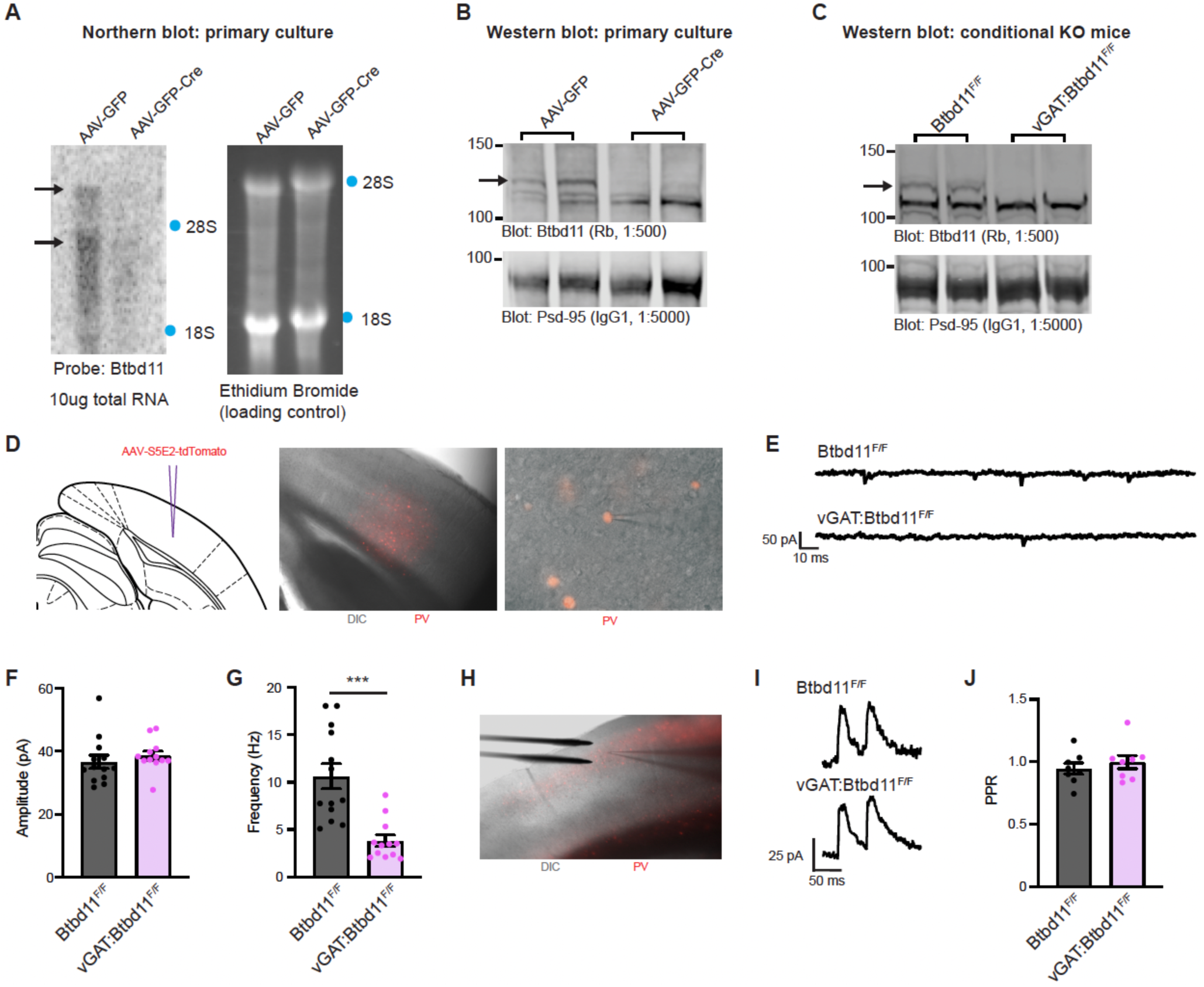
Btbd11 KO reduces glutamatergic signaling in PV-INs. (A) Northern blot to evaluate levels of Btbd11 mRNA in Btbd11^F/F^ cultures transduced with AAV-GFP (left lane) or AAV-GFP-Cre (right lane) and harvested at DIV14. 10μg of RNA was loaded onto a gel and ethidium bromide staining was used to confirm equal loading of RNA. Locations of 18S and 28S ribosomal RNA are indicated with blue dots. (B) Western blots characterizing the levels of Btbd11 (top blot) and Psd-95 (lower blot, as a loading control) in primary cortical Btbd11^F/F^ cultures transduced with AAV-GFP (left lanes) or AAV-GFP-Cre (right lanes). 12μg of protein was loaded from the PSD fraction (where Btbd11 is enriched). Cells were harvested at DIV12 and DIV14 (one sample per condition per timepoint). A black arrow indicates the band corresponding to Btbd11. (C) Western blots evaluating the levels of Btbd11 (top blot) and Psd-95 (lower blot, as a loading control) from the hippocampal PSD fraction of female control Btbd11^F/F^ animals or vGAT:Btbd11^F/F^ mice aged 3 months where Btbd11 is conditionally knocked out from INs. 20μg of protein was loaded. A black arrow indicates the band corresponding to Btbd11. (D) Schematic depicting the site of AAV-S5E2-tdTomato injection, used to visualize PV-INs in the V1 (left), with zoomed out (middle) and zoomed-in (right) merged image of DIC and mCherry fluorescence. (E) Example mEPSC traces recorded from V1 PV-INs in Btbd11^F/F^ or vGAT:Btbd11^F/F^ mice. (F and G) The mEPSC amplitude and frequency, respectively, of mEPSCs recorded from PV-INs from Btbd11^F/F^(black) animals or vGAT:Btbd11^F/F^ (magenta) mice. Error bars display S.E.M. *** indicates p < 0.001. (H) DIC image showing the placement of the electrical stimulating electrode for paired pulse ratio (PPR) recordings. (I) Example traces used to calculate the PPR in Btbd11^F/F^ or vGAT:Btbd11^F/F^ mice. (J) PPR data recorded from PV-INs from Btbd11^F/F^ (Black) or vGAT:Btbd11^F/F^ (magenta) mice. Error bars display S.E.M.

To create IN-specific Btbd11 knockout mice we bred vGAT^Cre/Wt^ mice with Btbd11^F/F^ animals to generate litters of vGAT^Wt/Wt^:: Btbd11^F/F^ (CON) and vGAT^Cre/Wt^::Btbd11^F/F^ (vGAT-Btbd11 KO) animals. Confirming our previous data that Btbd11 was specific to INs, we observed a total loss of Btbd11 in the hippocampal PSD fraction of vGAT-Btbd11 KO mice (**Figure 5C**). As PV-INs displayed the highest level of Btbd11 expression (based on RNA datasets, see **Figure 1D**) we decided to assess glutamatergic synapse function within PV-INs from CON or vGAT-Btbd11 KO mice by measuring miniature excitatory post synaptic potentials (mEPSCs) with whole-cell patch clamp recordings. We expressed tdTomato in the visual cortex PV-INs in CON or vGAT-Btbd11 KO littermates using AAVs with PV-specific enhancers (Vormstein-Schneider et al., 2020) to enable fluorescently guided recordings (**Figure 5D**,**E**). While the amplitude remained unchanged between the conditions (**Figure 5F**), we observed a dramatic decrease in the frequency of mEPSCs (**Figure 5G**). A frequency decrease could reflect fewer glutamatergic synapses in PV-INs of vGAT-Btbd11 KO mice, or a decreased presynaptic release probability. We measured the paired-pulse ratio (PPR) using electrical stimulation to explore presynaptic release probabilities (**Figure 5H**,**I**). The PPR was comparable between groups, suggesting that presynaptic release probabilities were not changed with Btbd11 knockout from INs (**Figure 5J**). Together, these data show that Btbd11^F/F^ mice are an effective tool to study Btbd11 KO and confirm the IN-specific nature of Btbd11 expression. Furthermore, we find that Btbd11 KO leads to decreased glutamatergic recruitment of PV-INs, likely though a postsynaptic mechanism.

### Loss of Btbd11 impacts circuit function *in vitro* and *in vivo*

INs are well known to play a critical role in regulating the activity of neuronal circuits. Having found that vGAT-Btbd11 KO mice have reduced glutamatergic recruitment of PV-INs, we speculated that network properties might be abnormal following Btbd11 KO. To test this, we first returned to our *in vitro* cell culture system and prepared primary hippocampal cultures from P0 Btbd11^F/F^ pups. We delivered GFP (CON) or GFP-Cre (KO) and jRGEGCO1 to cultures through AAV-transduction at DIV1-2 (**Figure 6A**). Spontaneous activity of the cultures was measured with live-cell imaging of the jRGECO1 signal to track Ca^2+^ dynamics. Large, and synchronous, Ca^2+^ transients were observed throughout the cultures (**Video S4**,**5**). We quantify these activity patterns in CON and KO cultures (**Figure 6B**). As expected for a dis-inhibited network, there was an increased frequency of Ca^2+^ transients in KO cultures (**Figure 6C**). Interestingly, the expression of PV protein was dramatically upregulated both in KO hippocampal cultures (observed with immunofluorescence; **Figure 6D**,**E**) and KO cortical cultures (observed with western blots; **Figure 6F**,**G**), potentially reflecting elevated activity since PV expression is regulated by activity of PV neurons (Donato et al., 2013).

**Figure 6.**
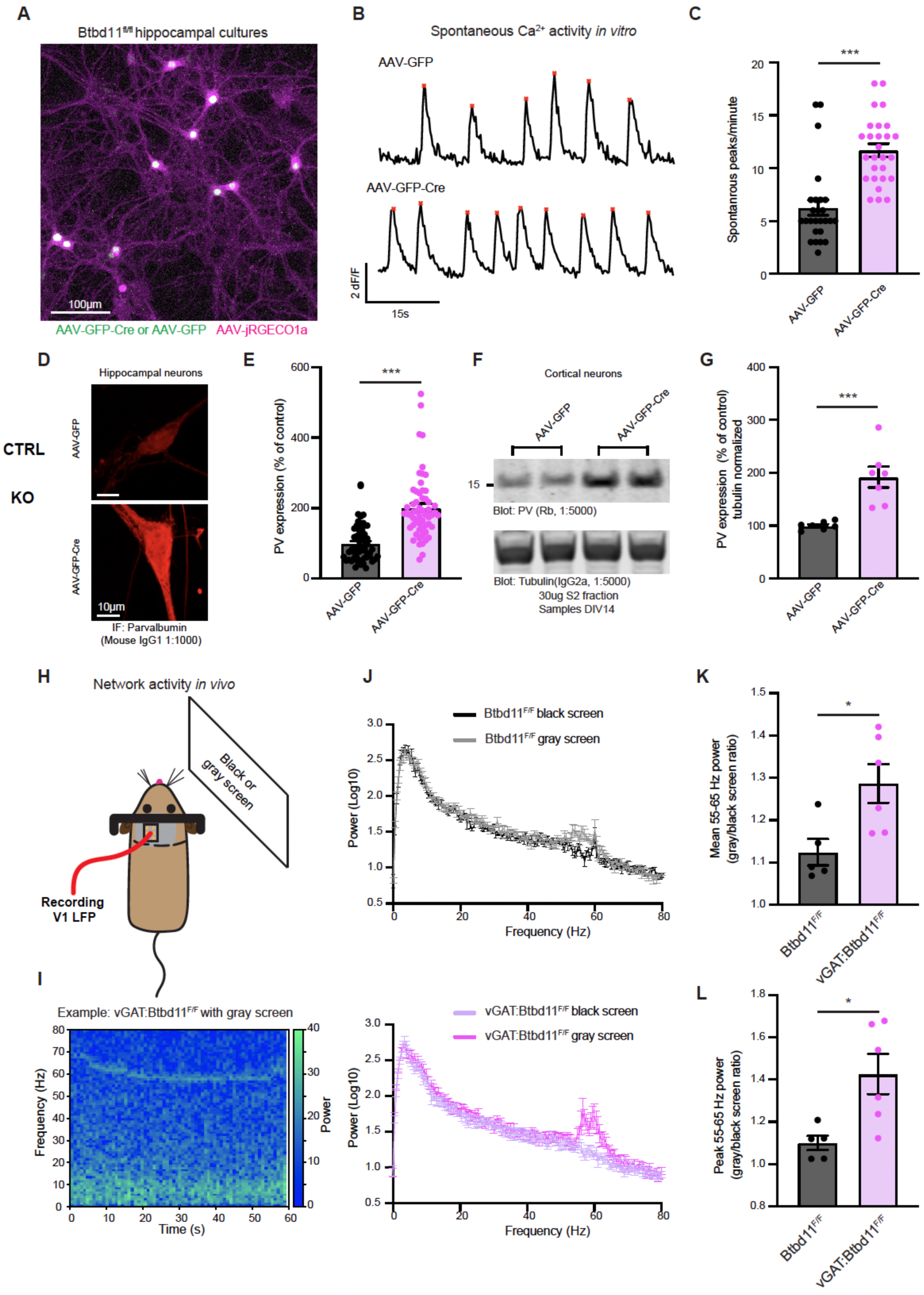
Loss of Btbd11 impacts circuit function *in vitro* and *in vivo*. (A) Confocal live-cell image of primary cultured hippocampal neurons from Btbd11^F/F^ mice transduced with AAV-jRGECO1a (magenta) and AAV-GFP or AAV-GFP-Cre (green). The example image is with AAV-GFP-Cre. Scale bar = 100μm. (B) Example traces for control (AAV-GFP, top) and knockout (AAV-GFP-Cre, bottom) cultures showing the average Ca^2+^ activity across multiple neurons in one field of view over a 60s period. Red stars indicate automatically identified peaks. (C) Quantification of large Ca^2+^ transients across multiple regions of interest and coverslips from 3 independent batches of neurons. Control (AAV-GFP) data is shown in black and knockout (AAV-GFP-Cre) data in magenta. *** indicates p <0.001. Error bars indicate S.E.M. (D) Confocal image showing immunofluorescence of PV in DIV14 primary cultures hippocampal Btbd11^F/F^ neurons transduced with AAV-GFP (control, top) or AAV-GFP-Cre (knockout, bottom). Scale bar = 10μm. (E) Quantification of PV immunofluorescence data with control data in black and knockout data in magenta. Error bars display S.E.M. (F) Western blot from the cytosolic S2 fraction of DIV14 primary cultured cortical Btbd11^F/F^ neurons transduced with AAV-GFP (control) or AAV-GFP-Cre (knockout). Top blot shows levels of PV, and the bottom blot shows alpha-tubulin used as a loading control. 30μg of lysate was run. (G) Quantification of western blot data evaluating levels of PV (normalized to alpha-tubulin levels) with control data in black and knockout data in magenta. Error bars display S.E.M. (H) Schematic of in vivo setup used to assess narrowband gamma oscillations in the V1 with presentation of a gray screen. (I) Example spectrogram showing the power over time in the 0-80 Hz range from the V1 of a vGAT:Btbd11^F/F^ mouse presented with a gray screen. Note the pronounced activity in the 55-65 Hz range. (J) Power spectra for Btbd11^F/F^ mice (n = 5, top; dark gray = dark screen, light gray = gray screen) and vGAT:Btbd11^F/F^ mice (n = 6, bottom; light magenta = dark screen, dark magenta gray = gray screen). Error bars indicate the S.E.M. (K and L) Quantification of the mean or peak 55-65 Hz activity, respectively, presented as a ratio of gray screen/black screen. Error bars display S.E.M. * indicates p < 0.05.

Next, we assessed circuit function *in vivo* using CON and vGAT-Btbd11 KO mice. PV-IN function has been closely tied to the regulation of fast local field potential (LFP) oscillations in the 30-100Hz “gamma” frequency (Cardin et al., 2009; Mann et al., 2005; Sohal et al., 2009). A narrowband gamma oscillation ∼55-65Hz can be induced in the visual cortex through presentation of a gray screen alone (Saleem et al., 2017). We exploited this straightforward assay to explore visually evoked gamma oscillations in the V1 of CON and vGAT-Btbd11 KO animals (**Figure 6H**,**I**). We implanted a tungsten electrode targeted to layer 4 of V1 as in (Cooke et al., 2015), and attached a metal bar to enable head-fixation of animals. After recovery from surgery, mice were handled and habituated to head restraint, then presented with either a black or gray screen (**Figure 6J**). As previously reported, we observed a narrowband gamma oscillation in the 55-65Hz range. Interestingly, we found that the gray/black screen ratio of both mean (**Figure 6K**) and peak (**Figure 6L**) 55-65 Hz activity was elevated in vGAT-Btbd11 KO animals compared to CONs indicating increased power in the narrowband gamma frequency. These data, *in vitro* and *in vivo*, argue that Btbd11 loss-of-function can impact the activity patterns of neuronal circuits, with important implications for their function in different states.

### Btbd11 KO mice are sensitized to challenge with an NMDA receptor antagonist

Having observed that Btbd11 knockout can impact circuit activity, we wondered if vGAT-Btbd11 KO mice impacts animal behavior. We assessed the behavior of CON and vGAT-Btbd11 KO animals in an open field to assess locomotor activity. Animals were run in the dark, with infrared beam breaks used as a readout of activity (**Figure 7A**). No differences were observed in terms of the total beam breaks (**Figure 7B**). We subsequently assessed short-term spatial memory using a spontaneous alternation version of the Y-maze. We observed no difference between CON and vGAT-Btbd11 KO animals, indicating that under basal conditions spatial short-term memory was intact (**Figure 7C**).

**Figure 7.**
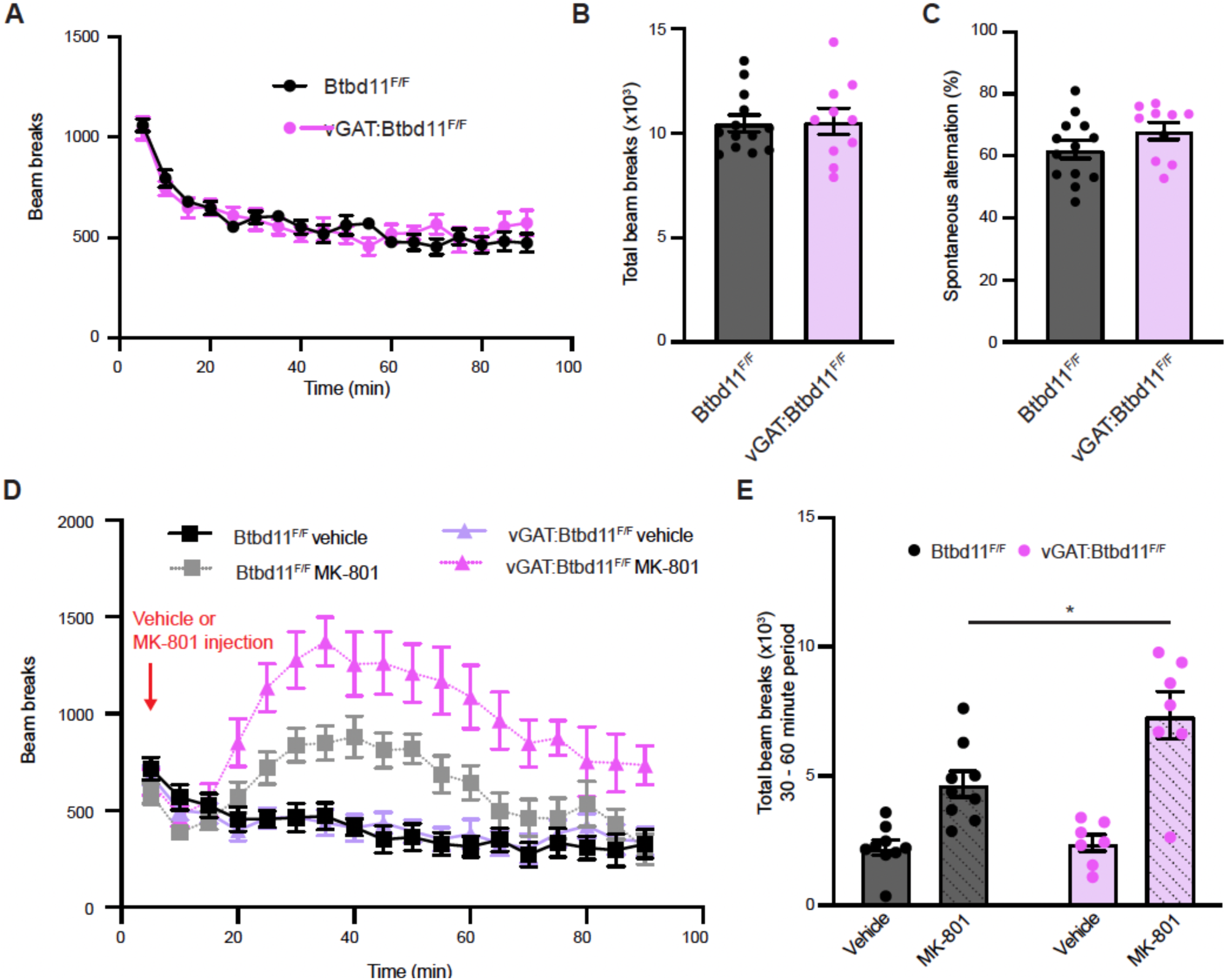
Btbd11 KO mice are sensitized to challenge with an NMDA receptor antagonist. (A) Locomotor activity of Btbd11^F/F^ (black) and vGAT:Btbd11^F/F^ (magenta) mice exploring a novel environment. Infrared beam breaks were used as a proxy of locomotor activity. Error bars indicate S.E.M. (B) Total beam breaks in the 90 min exploration period for Btbd11^F/F^ (black) and vGAT:Btbd11^F/F^ (magenta) animals. Error bars indicate S.E.M. (C) Spontaneous alternation of Btbd11^F/F^ (black) and vGAT:Btbd11^F/F^ (magenta) animals in a Y-maze test of short-term spatial memory. Error bars indicate S.E.M. (D) Locomotor activity of Btbd11^F/F^ (black and gray) and vGAT:Btbd11^F/F^ (light and dark magenta) mice in an open field arena following injection with either saline or MK-801 (0.2mg/kg). Infrared beam breaks were used as a proxy of locomotor activity. Error bars indicate S.E.M. (E) Quantification of the total infrared beam breaks in the 30-60-min period after injection in Btbd11^F/F^ (black) and vGAT:Btbd11^F/F^ (magenta) mice. * Indicates p < 0.05.

Genetic manipulations that impact glutamatergic synapses in INs often display altered sensitivity to NMDA receptor antagonist challenge. For example, mice which lack NMDARs in PV-INs are more sensitive to the effects of the NMDA receptor antagonist MK-801 (Bygrave et al., 2016). Since vGAT-Btbd11-KO mice had reduced glutamatergic recruitment of PV-INs we speculated that they might be predisposed to MK-801 challenge. Therefore, we administered CON and vGAT-Btbd11 KO mice with MK-801 (0.2 mg kg^-1^) or saline (as a vehicle) and measured their locomotor response in the open field apparatus, this time in the light (**Figure 7D**). MK-801 administration led to a dramatic increase in locomotor activity, however, this was particularly prominent in vGAT-Btbd11 KO mice, indicating a increased sensitivity to the drug (**Figure 7E**). These data show that the underlying circuitry in mice which lack Btbd11 in INs sensitizes them to subsequent NMDA receptor challenge.

## DISCUSSION

In this study we identify Btbd11 as a novel inPSD protein and characterize its basic properties and function using biochemistry, imaging, electrophysiology, and behavior.

### Glutamatergic PSDs show cell-type-specific protein specializations

Using immunoisolation and proteomics we demonstrate that it is possible to identify inPSD specific proteins. To the best of our knowledge, this is the first attempt to uncover differences in PSD composition between INs and ENs using proteomics. A similar method was used to examine differences in PSDs from different subpopulations of ENs (Zhu et al., 2020), highlighting the usefulness of the approach. Initially we focused our effort into a thorough characterizing of Btbd11. In future studies it will be interesting to explore the other putative inPSD proteins (**Table S1**). Furthermore, by combining proteomic with new methods such as *in vivo* proximity labeling (Branon et al., 2018; Uezu et al., 2016), it should be possible to uncover yet more PSD protein specializations, including those that do not form complexes with Psd-95 which would likely be missed by our screen.

### Btbd11 is localized exclusively to inPSDs in cortex and hippocampus

Converging evidence shows that Btbd11 is expressed exclusively in INs within cortical and hippocampal tissue, however, it is possible that in other brain regions this cell-type-specificity will be lost. It is noteworthy that Btbd11 was previously identified as a reliable marker of cortical interneurons, although no function or properties of Btbd11 were explored (Rossier et al., 2015). The subcellular distribution of Btbd11 is highly targeted to glutamatergic synapses—via interactions with Btbd11’s PBM. Our yeast 2-hybrid screen indicates putative interacting proteins, including known synaptic proteins that impact synapse function. It will be important to further characterize the interactome of Btbd11 to gain further mechanistic insights into how it regulates glutamatergic synapses in INs.

### LLPS of Btbd11 with Psd-95

We show that Btbd11 undergoes LLPS with Psd-95. LLPS is highly sensitive to protein concentration, but since the PSD concentrates synaptic proteins, we expect the concentration of Btbd11 and Psd-95 to surpass a threshold to support biomolecular condensation at glutamatergic synapses in INs. Thus far, difficulties in purifying full-length Btbd11 (which we find is necessary for LLPS properties) have prevented *in vitro* studies to calculate the precise concentration of Btbd11 and Psd-95 required to trigger LLPS. Phase separation at the PSD is likely ubiquitous among neuronal cell-types and has been argued to underscore the assembly and stability of the PSD (Chen et al., 2020; Feng et al., 2019; Zeng et al., 2016). It is tempting to speculate that Btbd11 can stabilize Psd-95 at the synapse through promoting LLPS. If this was indeed the case it likely reflects a generalizable phenomenon, for which we have identified a particular cell-type-specific specialization involving Btbd11 utilized by INs. LLPS of the iPSD may play an even more critical role than it does at the ePSD as the iPSD is exposed on the dendritic shaft and is not compartmentalized in spine-like protrusions thought to limit diffusion to and from the synapse.

### Reduced glutamatergic recruitment in PV neurons of Btbd11 KO mice

With whole-cell patch clamp recordings we observed a decrease in the mEPSC frequency in PV-INs when Btbd11 was knocked out from INs. This phenotype is similar to that of IN-specific deletion of ErbB4 or global knockout of Brevican, two proteins that are also enriched at glutamatergic synapses in PV-INs (Favuzzi et al., 2017; del Pino et al., 2013). Because a measure of presynaptic release (PPR) was unchanged with Btbd11 knockout, we expect that this mEPSC phenotype is a result of decreased glutamatergic synapses within PV-INs. We expect this is a consequence of a destabilizing effect on Psd-95 due to Btbd11 knockout, as we found that Btbd11 overexpression was able to stabilize Psd-95 at the PSD of cultured INs. It will be interesting to test if other IN subtypes, such as those expressing somatostatin, also receive reduced glutamatergic input. Furthermore, in subsequent studies it will be interesting to elucidate if there is a time dependence of Btbd11 deletion, or if this phenotype is dependent on gene deletion early in development (Cre switches on early in the vGAT^Cre^ line).

### Altered network properties and sensitivity to NMDA receptor challenge in Btbd11 KO mice

In cultured Btbd11 knockout neurons we observed an increase in spontaneous Ca^2+^ transients, consistent with a lack of inhibition in the cultures. Furthermore, we observed an exaggerated induced gamma frequency oscillation *in vivo* when mice were presented with a gray screen. Because of the role PV-INs have in regulating the firing of other neurons and supporting synchronous activity, we suspect that altered activity of PV-INs—through loss of glutamatergic signaling—is responsible for these changes in network properties. Behaviorally, we observed that IN-specific Btbd11 knockout mice are sensitized to NMDA receptor antagonism with MK-801. Previous work has shown that INs could be preferentially sensitive to NMDA receptor antagonism at certain doses (Homayoun and Moghaddam, 2007). Furthermore, deletion of NMDA receptors from different populations of INs results in differential sensitivity to MK-801 challenge (Belforte et al., 2010; Bygrave et al., 2016, 2019; Cardin et al., 2009). The increased sensitivity vGAT-Btbd11 KO mice to MK-801 likely manifests because of their decreased number of glutamatergic synapses in PV-INs. In future studies it will be interesting to test if other stressors, such as post-weaning isolation (Belforte et al., 2010; Jiang et al., 2013) or reduced environmental enrichment (Bygrave et al., 2019) have exaggerated effects on vGAT-Btbd11 KO mice compared to CON animals.

In summary, we reveal that Btbd11 as an IN-specific glutamatergic synaptic protein and show that it plays an important role in regulating glutamatergic synapses in INs using biochemistry and *in vitro* live-cell imaging experiments through to *in vivo* physiology and behavior. It will be important to uncover if Btbd11—or other inPSD proteins—could be potential therapeutic targets for neurological disorders in which GABAergic signaling is disrupted.

## ACKNOWLEDGEMENTS

We would like to thank members of the Huganir laboratory for helpful discussion. Alexei Bygrave was supported by a K99/R00 award (K99MH124920). R01NS036715 awarded to Richard Huganir also supported this work and RF1MH120119 to Haining Zhong. We would like to thank Sarah Rodriguez for help with mouse colony management, Chip Hawkins for conducting IVF to generate Btbd11 mice, Ingie Hong and Elena Lopez-Ortega for help with in vivo electrophysiology recordings and analysis pipelines, William Hale for help with primary cultured mouse neurons, Mike Delannoy for processing of samples for electron microscopy, Paul Worley for useful discussions and input on experimental design, and Tatiana Boronina and Robert Cole for running and analyzing mass spectrometry data.

## SUPPLEMENTARY FIGURES

**Figure S1.**
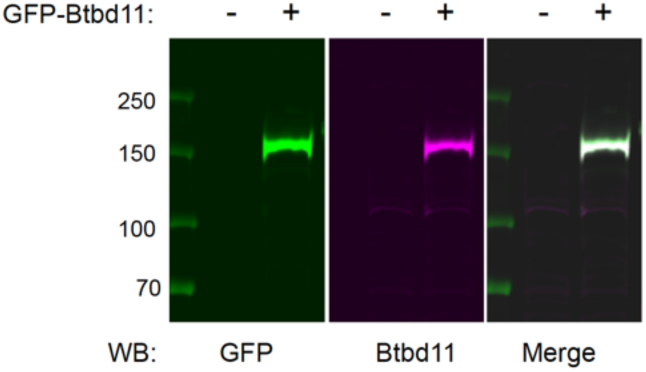
Validation of Btbd11 antibody, *related to Figure 1* Western blot using lysate from HEK cell transfected with GFP-Btbd11 or non-transfected cells. Blots were probed with antibodies against Btbd11 (magenta) or GFP (green). The merged image shows the overlap of the bands identified with each antibody.

**Figure S2.**
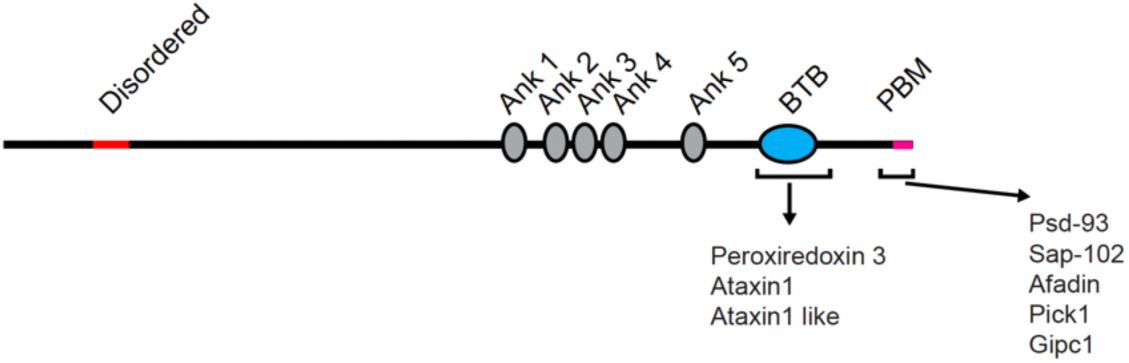
Yeast 2-hybrid screen identifies putative Btbd11 interaction partners, *related to Figure 2* Schematic depiction of results from the yeast two-hybrid screen with the BTB domain or C-terminal portion of Btbd11 containing the PBM.

**Figure S3.**
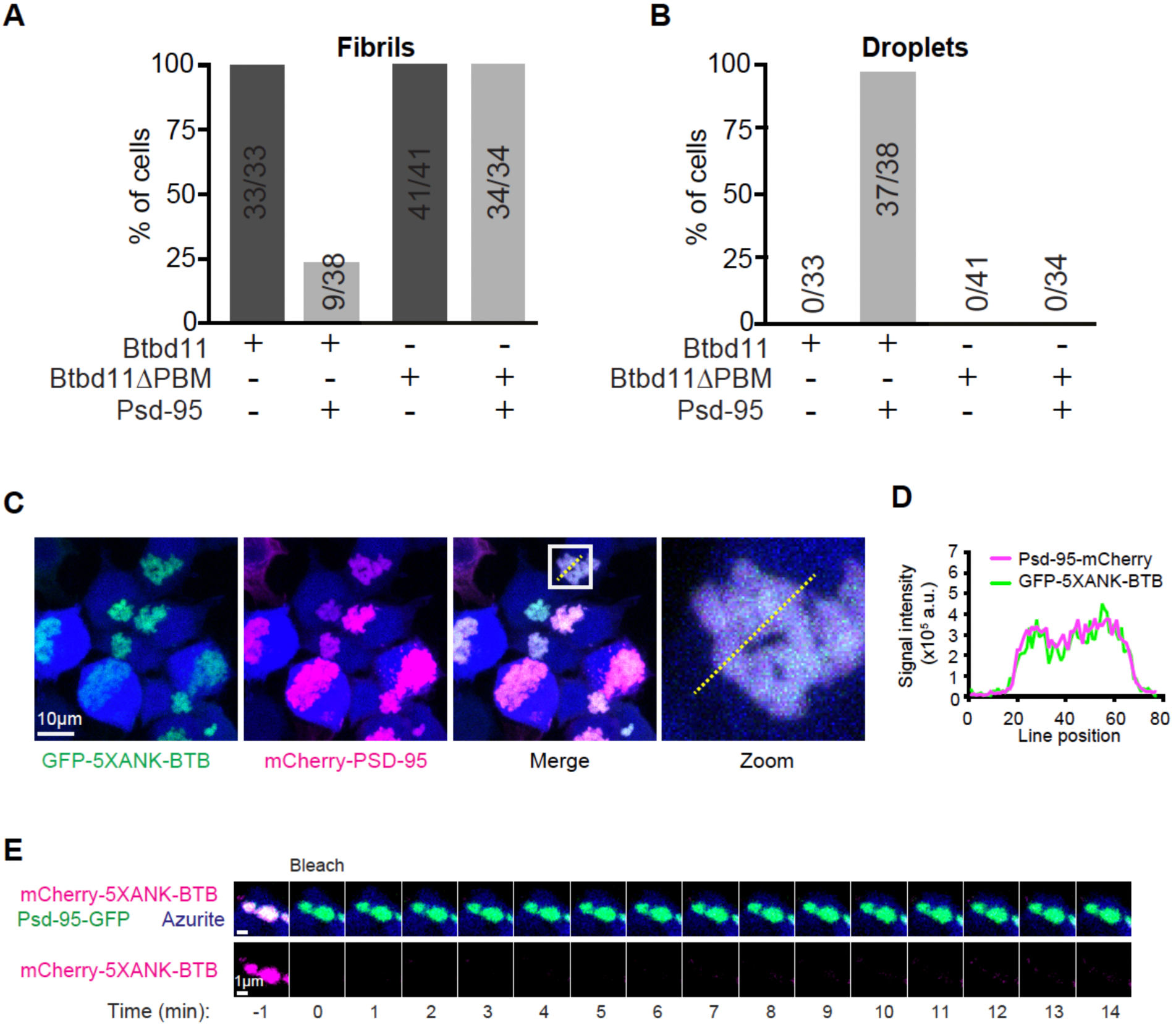
Liquid-liquid phase separation of Btbd11, *related to Figure 3* (A and B) Proportion of HEK cells which contain fibril-like assemblies or droplets, respectively, when GFP-Btbd11 or GFP-Btbd11ΔPBM is expressed with or without Psd-95-mCherry. (C) Confocal image of HEK cells expressing GFP-5XANK-BTB (green), Psd-95-mCherry (magenta) and azurite (blue). Scale bar indicates 10μm. The white boxed area is enlarged on the right. The yellow dotted line indicates where a line-scan was evaluated. (D) Line scan showing signal intensity of GFP-5XANK-BTB (green) and Psd-95-mCherry (magenta). (E) FRAP of a mCherry-5XANK-BTB (magenta) and Psd-95-GFP (green) puncta with a composite image shown in the upper panels and just mCherry-5XANK-BTB shown on the bottom. Scale bar = 1μm. Note the lack of mCherry-5XANK-BTB signal recovery.

**Figure S4.**
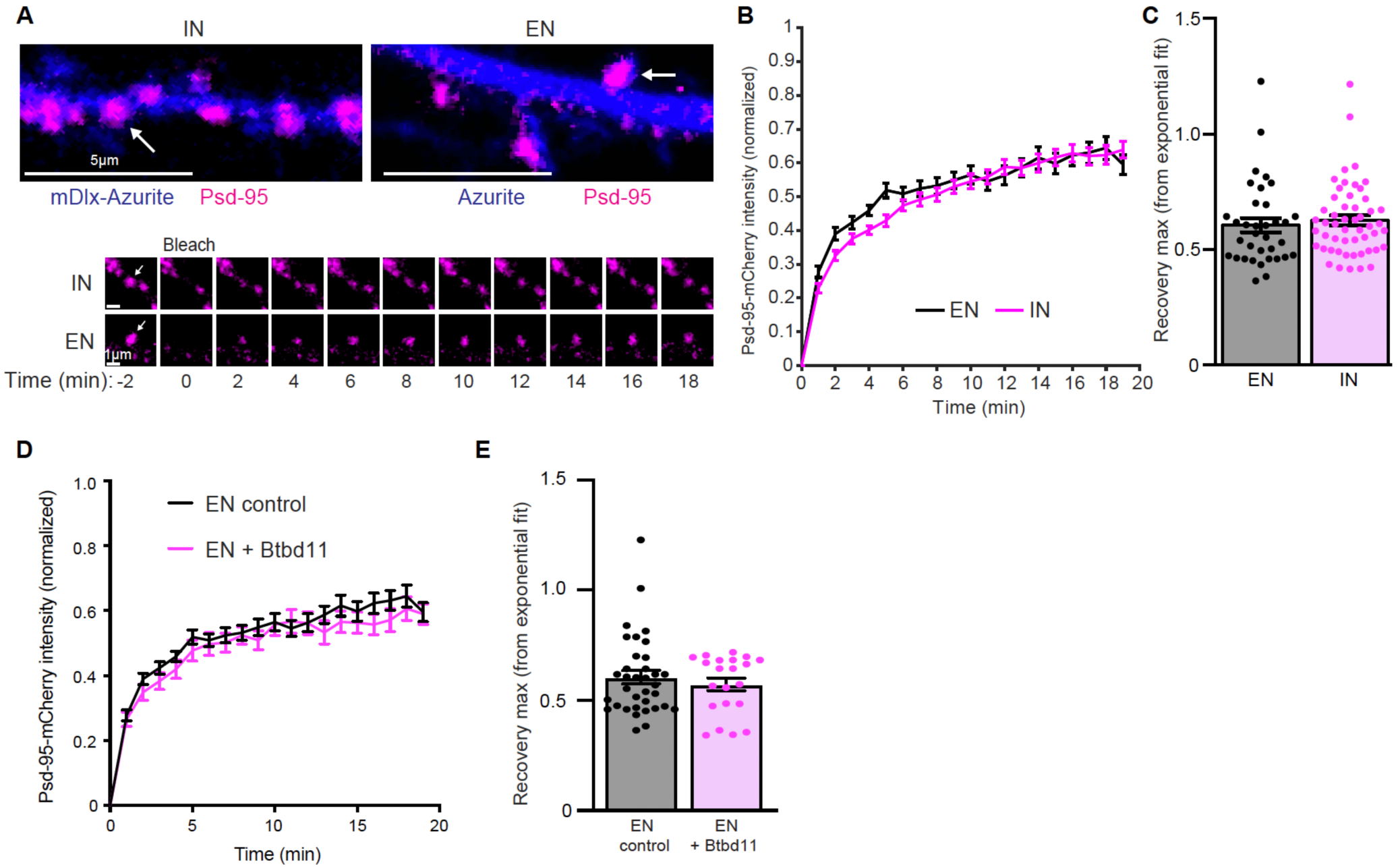
FRAP of Psd-95 in excitatory neurons, *related to Figure 4* (A) Live-cell confocal imaging and FRAP of Psd-95-mCherry (magenta) in a putative interneuron (IN, identified with mDlx-Azurite) or excitatory neuron (EN, identified by the presence of dendritic spines with an Azurite cell-fill). The lower panels show FRAP of the individual puncta labeled in the upper panels with a white arrow. Respective scale bars are indicted at the bottom left of each image. (B) Quantification of Psd-95-mCherry FRAP in ENs (black) or INs (magenta). Error bars display S.E.M. (C) Quantification of the estimated recovery maximum from an exponential fit of the FRAP data for each bleached punctum. EN data is shown in black and IN data is in magenta. Error bars display S.E.M. (D) Quantification of Psd-95-mCherry FRAP in ENs alone (black) or with overexpression of Btbd11 (magenta). Error bars display S.E.M. (E) Quantification of the estimated recovery maximum from an exponential fit of the FRAP data for each bleached punctum. EN control data is shown in black and EN data with Btbd11 overexpression is in magenta. Error bars display S.E.M. Note the EN data without Btbd11 in panel (B) is the same as the control data presented in panel (D).

**Figure S5.**
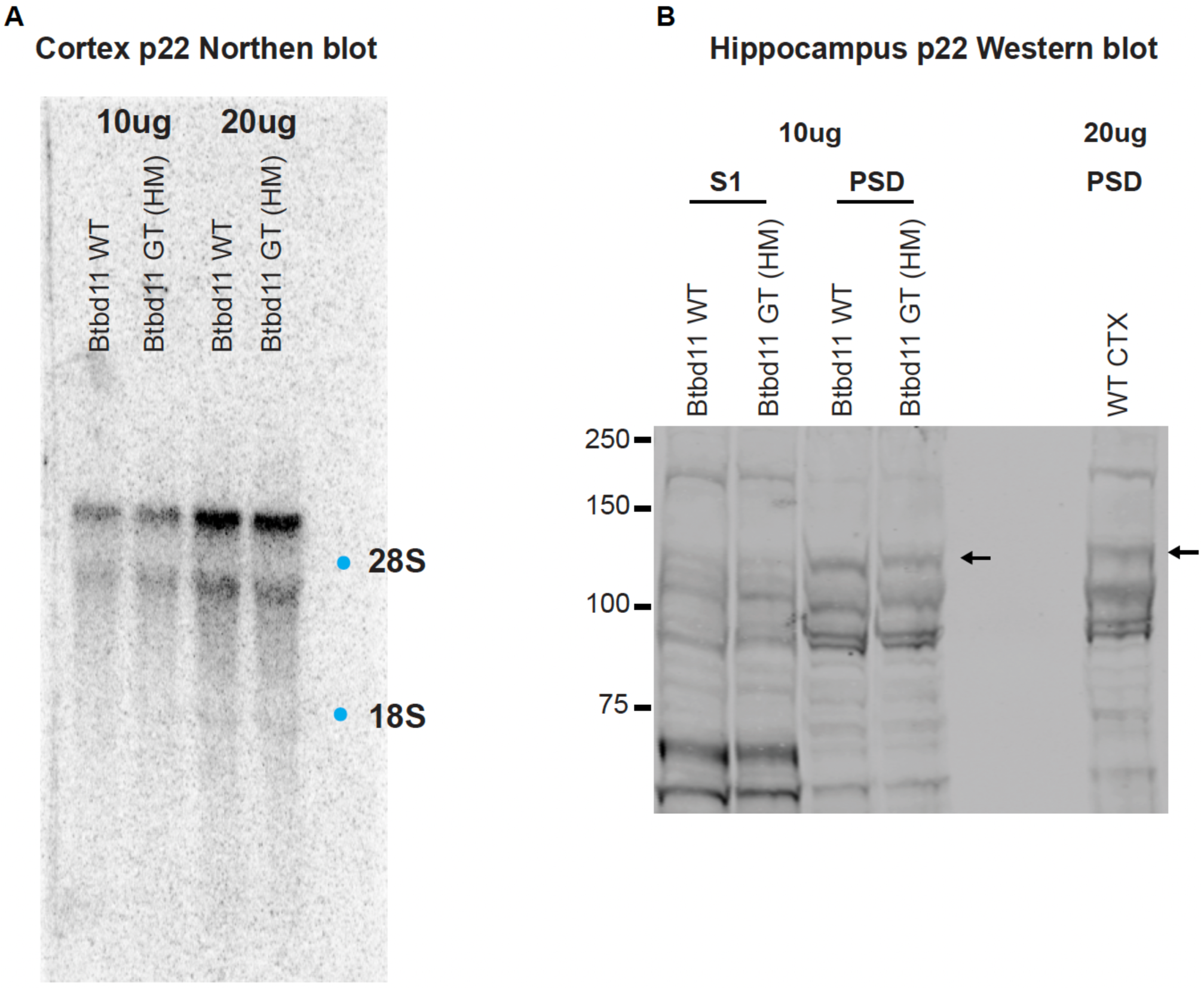
Characterization of Btbd11 Gene Trap mice, *related to Figure 5* (A) Northern blots to assess the levels of Btbd11 mRNA in the cortex of postnatal day 22 Btbd11 wildtype (WT) or homozygous Btbd11 Gene Trap (GT) animals. Either 10μg or 20μg of mRNA was loaded. A blue dot indicates the location of 18S and 28S ribosomal RNA. (B) Western blots to assess the level of Btbd11 in the hippocampus of Btbd11 wildtype (WT) or homozygous Btbd11 Gene Trap (GT) animals. 10μg of the PSD fraction was run, and membranes were probed with an antibody for Btbd11. A black arrow indicates the band corresponding to Btbd11. On the right 20μg of WT cortical PSD fraction was run to ensure that endogenous Btbd11 could be detected.

**Figure S6.**
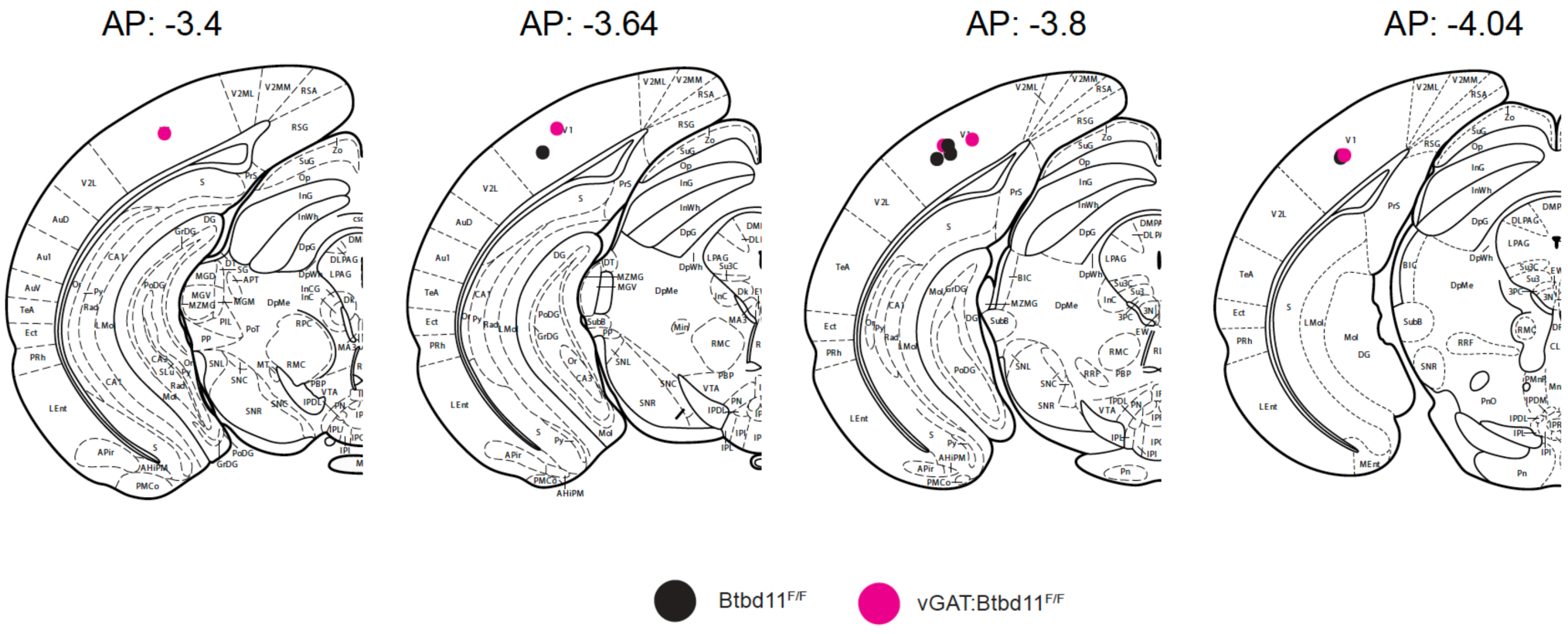
Electrode placement reconstructions, *related to Figure 6* Reconstructions of estimated electrode positioning based on electrolytic lesions. Black dots = Btbd11^F/F^ and magenta dots = vGAT:Btbd11^F/F^ animals, respectively. AP = anterior/posterior location relative to bregma. Note: there are two vGAT:Btbd11^F/F^ with overlapping estimated lesion sites in the AP: -4.04 image.

## SUPPLEMENTARY TABLES

**Table S1.** Mass spectrometry results, *related to Figure 1* Table of inPSD candidates identified in the cell type-specific Psd-95-GFP immunoisolation experiment. Proteins were quantified with label-free quantification and then normalized to levels of Psd-95. Then the vGAT/CaMKII ratio was calculated to identify putative proteins enriched at the PSD of inhibitory interneurons.

| inPSD hits with vGAT/CaMKII ratio with value > mean + 2x standard deviation |  |  |
| --- | --- | --- |
| Accession | Description | log norm ratio (vGAT/CaMKII) |
| NP_082985.2 | ankyrin repeat and BTB/POZ domain-containing protein BTBD11 isoform 1 [Mus musculus] | 3.37 |
| NP_058017.1 | stress-induced-phosphoprotein 1 [Mus musculus] | 2.41 |
| XP_006534552.1 | PREDICTED: myosin-10 isoform X1 [Mus musculus] | 2.23 |
| NP_076288.1 | protein O-GlcNAcase [Mus musculus] | 2.08 |
| XP_006515861.1 | PREDICTED: TNF receptor-associated factor 3 isoform X1 [Mus musculus] | 2.05 |
| XP_006498025.1 | PREDICTED: E3 ubiquitin-protein ligase LRSAM1 isoform X1 [Mus musculus] | 2.02 |
| XP_006538016.1 | PREDICTED: F-box/LRR-repeat protein 4 isoform X1 [Mus musculus] | 2.00 |
| XP_017175463.1 | PREDICTED: fatty-acid amide hydrolase 1 isoform X1 [Mus musculus] | 1.93 |
| XP_006527892.1 | PREDICTED: glycerol kinase isoform X1 [Mus musculus] | 1.92 |
| NP_659136.1 | palmitoyltransferase ZDHHC5 [Mus musculus] | 1.89 |
| XP_017171979.1 | PREDICTED: oxidation resistance protein 1 isoform X1 [Mus musculus] | 1.88 |
| XP_006511788.1 | PREDICTED: E3 ubiquitin-protein ligase ARIH2 isoform X1 [Mus musculus] | 1.87 |
| NP_085041.1 | E3 ubiquitin-protein ligase RNF34 [Mus musculus] | 1.86 |
| XP_006519125.1 | PREDICTED: E3 ubiquitin-protein ligase RNF31 isoform X1 [Mus musculus] | 1.83 |
| XP_006537872.1 | PREDICTED: PHD finger protein 24 isoform X1 [Mus musculus] | 1.75 |
| NP_033750.1 | a disintegrin and metallopeptidase domain 4 precursor [Mus musculus] | 1.72 |
| XP_011249102.1 | PREDICTED: glutamate receptor ionotropic, NMDA 2D isoform X1 [Mus musculus] | 1.68 |
| XP_006533949.1 | PREDICTED: mitochondrial Rho GTPase 1 isoform X1 [Mus musculus] | 1.65 |
| NP_035234.1 | protein kinase C epsilon type [Mus musculus] | 1.64 |
| XP_006526184.1 | PREDICTED: E3 ubiquitin-protein ligase RNF14 isoform X1 [Mus musculus] | 1.62 |
| XP_006516266.1 | PREDICTED: WD repeat-containing protein 20 isoform X1 [Mus musculus] | 1.62 |
| NP_033534.2 | vesicular inhibitory amino acid transporter [Mus musculus] | 1.60 |

**Table S2.**
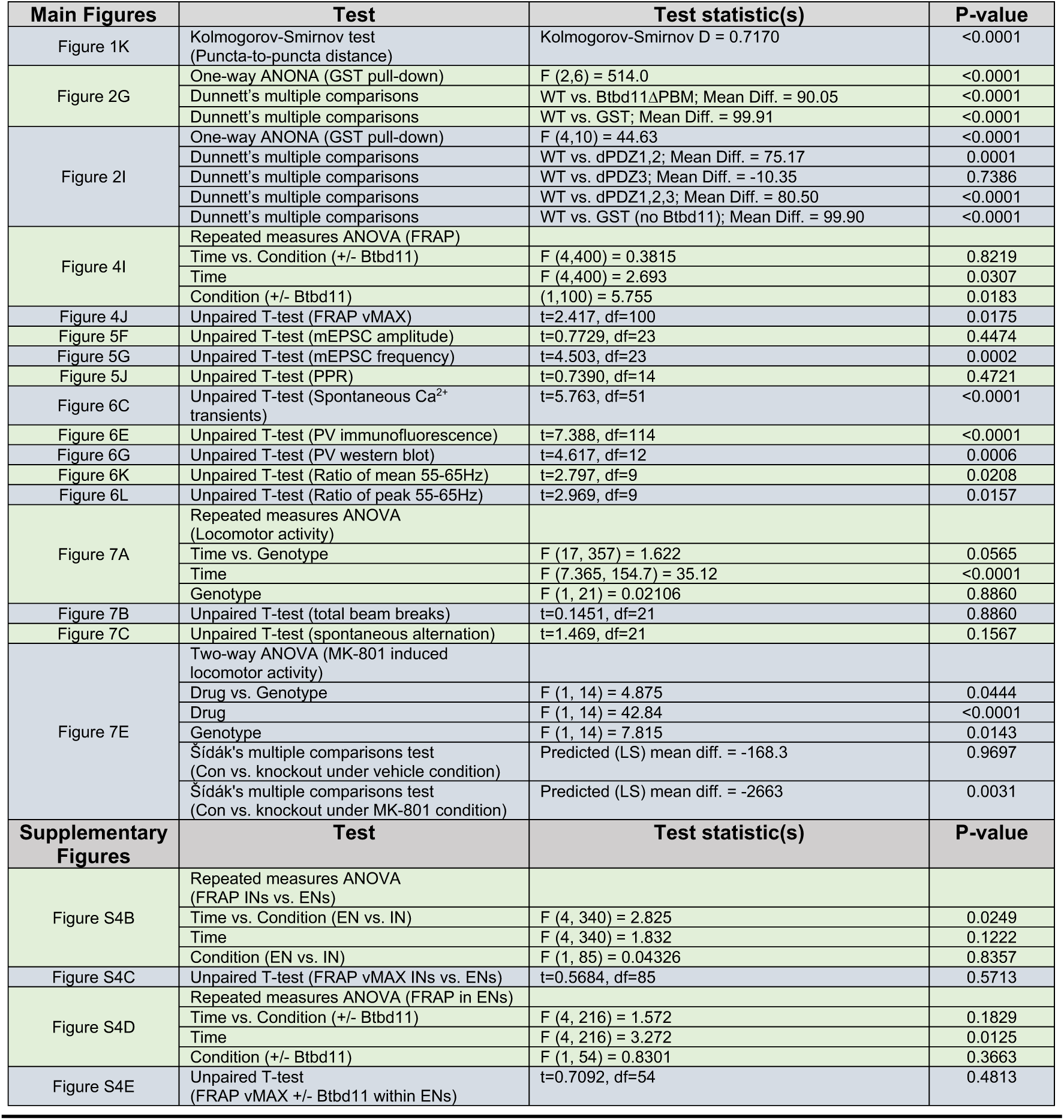
Details of statistical tests run, *related to Figures 2 and 4-7* Table with statistical details for all tests performed.

## VIDEO TITLES

**Video S1**

Alpha Fold prediction of Btbd11 structure. Based on UniProt annotations and the recognition of a putative PDZ binding motif (PBM) portions of the protein are shaded as follows: red = disordered, gray = ankyrin repeats (Ank), blue = BTB domain (BTB), magenta = PDM.

**Video S2**

Time-lapse of two GFP-Btbd11 (green) and Psd-95-mCherry (magenta) puncta fusing over time. The cell also contains an azurite cell-fill (blue). This is the same data presented in **Figure 3K**.

**Video S3**

Longitudinal live-cell imaging of a hippocampal neuron expressing an Azurite cell-fill (Blue) and GFP-Btbd11 (Green). A portion of a GFP-Btbd11 fibril is bleached. Note the lack of fluorescence recovery.

**Video S4**

Live cell imaging of jRGECO1a in control cultures (Btbd11^F/F^ with AAV-GFP). Note the synchronous Ca^2+^ transients.

**Video S5**

Live cell imaging of jRGECO1a in Btbd11 knockout cultures (Btbd11^F/F^ with AAV-GFP-Cre). Note that the synchronous Ca^2+^ transients appear more frequent than in control conditions (**Video S4**).

## METHODS

### Animal care

All animals were treated in accordance with the Johns Hopkins University Animal Care and Use Committee guidelines.

### GFP-immunoisolation

Cortex and hippocampus from 1 CaMKII:Psd-95-GFP and 4 vGAT:Psd-95-GFP animals were collected from male mice aged 11-12 weeks. Brain tissue was homogenized in 8ml of ice-cold lysis buffer (1% DOC, 50mM Tris (pH9), 50mM NaF, 20μM ZnCl_2_, 1mM Na_3_VO_4_) with the addition of Pefabloc SC (Roche), Okadaic acid (200nM) and a protease inhibitor cocktail (Roche Complete) with a Dounce homogenizer and solubilized for 1 hour rotating at 4°C. Samples were then clarified with spins at 17,000g and 50,000g each for 30min and at 4°C. Further lysis buffer was added so that GFP-immunoisolation experiments were set up in 10ml total volume. For each sample 50μl of pre-washed GFP-TRAP agarose beads (Chromotek) was added and incubated with rotation overnight at 4°C. Beads were washed x6 with ice-cold lysis buffer (with same inhibitors as before) using spin-filter columns (spin 1500g for 1 minute). Samples were eluted with 1% SDS containing 2.5% beta-mercaptoethanol. The vGAT samples were serially eluted to increase the concentration of eluant (since the abundance of Psd-95-GFP is much lower in the vGAT:Psd-95-GFP mice). Samples were then subject to mass spectrometry.

Note: the same pull-down procedure was followed to get samples for western blots (**Figure 1C**), except 5x vGAT:Psd95-GFP mice were pooled.

### Mass spectrometry

GFP-immunoisolation samples (see above) were reduced with DTT, alkylated with IAA and TCA/Acetone precipitated with x8 volume overnight at 37°C. Samples were then re-constituted in 12uL 500mM TEAB, 48uL water and 25ng/uL trypsin/LysC added. Samples were proteolyzed at 37°C overnight. Peptides were desalted on Qasis U-HLB plates, eluted with 60% acetonitrile/0.1%TFA, and dried. Desalted peptides, 10% and 20% were analyzed by nano-LC/MS/MS on QExactive Plus at resolution 140K on precursor and 35K on fragment, using 85min gradient from 2% acetonitrile/0.1% formic acid to 98% acetonitrile/0.1% formic acid. Injected peptides showed abundant base peak chromatographs. Database search: FilesRC option was applied and MSMS spectra were searched with Mascot 6.1 via Proteome Discovere 2.3 against RefSeq2017_83_mouse and a small database with added enzymes, bovine serum albumin (our standard). Mass tolerance 4ppm for precursor, 0.01Da for fragment variable modifications: carbamidomethylation on Cys, oxidation on M, deamidation NQ. Peptides validated with Percolator. Mascot .dat files compiled in Scaffold. Note: just the 20% samples were used for analysis. Label-free quantification was performed, and protein abundances were normalized to the abundance of Psd-95. The ratio of vGAT/CaMKII abundance was then calculated to identify putative inPSD proteins (plotted in **Figure 1B**). From the ratio data, proteins with ratios greater than the mean + 2X the standard deviation were classified as putative inPSD candidates (**Table S1**).

### Western blots

Samples in Laemmli Buffer were heated to 60°C for 15 minutes prior to running western blots. Depending on target protein Mw, precast 4-12% or 8% Bis-Tris gels were used (Thermo Fisher Scientific) with NuPAGE MOPS SDS running buffer (Thermo Fisher Scientific). Proteins were transferred to nitrocellulose membrane (GE Healthcare) and blocked for 1 hour with Odyssey Blocking Buffer (Li-COR bioscience) before probing with primary antibodies (see below) overnight at 4°C or at room temperature for 1-2 hours. Antibodies were made up in TBS supplemented with 3% BSA and 0.1% Tween-20. Blots were washes with TBS with 0.1% Tween-20 (TBST) 4x and incubated with secondary antibodies (all fluorescently conjugated with 680 or 800nm dyes and used at 1:10000 and from Li-COR bioscience) made up in TBS supplemented with 3% BSA and 0.1% Tween-20 and 0.02% SDS for 1 hour at room temp. Blots were washed 4x in TBST and imaged on a Li-COR scanner. Bands were quantified in Image Studio (Li-COR bioscience). *Primary antibodies used:* Tanaka anti dsRED (Living colors, rabbit polyclonal, 1:1000), Santa Cruz anti -GFP (B2, IgG2_a_, 1:2000), homemade anti-Btbd11 (rabbit polyclonal, 1:500), Neuromab anti-Psd-95 (K28/74 mouse IgG_1_, 1:5000), Sigma anti-GST (GST-2, mouse IgG2_B_, 1:5000), Swant anti-Parvalbumin (PV27, rabbit polyclonal, 1:5000), Santa Cruz anti-alpha-tubulin (B-7, mouse IgG2_a,_ 1:5000).

### Immunofluorescence

Neurons were rinsed 1X with PBS at room temperature and then fixed for 15 minutes at room temperature with 4% paraformaldehyde (Electron Microscopy Sciences) made up in PBS and supplemented with 4% sucrose. For certain experiments (**Figure 1G**) this fixation step was replaced with a 20-minute fixation in methanol at -20°C. In other instances, to promote Gad-67 and Psd-95 immunofluorescence, cells were subject to a 90s methanol wash (−20°C) following the paraformaldehyde fixation described above. Cells were washed 4x with PBS and incubated in primary antibodies (see below) in a pH 7.4 gelatin buffer (30 mM phosphate buffer, 0.2% gelatin, 0.3% Triton X-100, and 0.25M NaCl) overnight at 4°C. Cells were washed 4x with PBS and then incubated with secondary antibodies in the same GDB buffer for 1 hour at room temperature. All 488, 568 and 647 secondary antibodies were raised in goat, Alexa Fluor conjugated, used at 1:500 and purchased from Thermo Fisher Scientific. All 405 conjugated antibodies were raised in goat, DyLight 405 conjugated, used at 1:500, and purchased from Jackson Immuno Research Laboratories. Cells were washed 4x in PBS, briefly rinsed in distilled water and mounted on slides with PermaFluor mounting media (Thermo Fisher Scientific). Slides were stored in the dark at 4°C. *Primary antibodies used:* Abcam anti-GFP (ab13970, chicken, 1:2000), Neuromab anti-Psd-95 (K28/43, mouse IgG2_a_, 1:500), homemade anti-Btbd11 (rabbit polyclonal, 1:100), Tanaka anti dsRED (Living colors, rabbit polyclonal, 1:1000), Santa Cruz anti-Gad-67 (F-6, mouse IgG_3_, 1:100), Swant anti-parvalbumin (Clone 235, mouse, IgG_1_, 1:1000), Synaptic Systems anti-Gephyrin (147 011, mouse, IgG_1_, 1:500).

### Yeast two-hybrid experiments

Different combinations of pDBLeu-Btbd11-C-terminus (consisting of the terminal 58 or 54 amino acids of Btbd11 for Btbd11 and Btbd11ΔPBM, respectively) and pPC86 -PSD95 PDZ constructs were co-transformed into PJ69 yeast cells and bait/prey pairs were selected for by growth on -Leu, Trp media at 30°C. Btbd11-PSD95 interactions were then tested by the ability of the different yeast bait/prey combinations to grow on -Leu, Trp, His media.

To identify putative Btbd11 interaction partners the BTB domain of Btbd11 or the C-terminal region of Btbd11 were cloned into pDBLeu and used as baits in yeast 2-hybrid screens with a rat hippocampal library. Plasmids from yeast clones that could grow on -Leu,Trp, His, Ade media were rescued and analyzed by sequencing.

### Cell-culture

#### HEK cells

HEK cells were grown on 10cm plates (for biochemistry) or on collagen (Advanced Biomatrix) coated glass coverslips in 12-well plates (for immunofluorescence and live-cell imaging) in DMEM (Gibco) supplemented with 10% Fetal Bovine Serum (Hyclone) and Penicillin-Streptomycin antibiotics (100 U/ml, Thermo Fisher Scientific). For biochemistry (GST-Pulldown experiments) HEK cells were transfected via Calcium Phosphate precipitation. For immunofluorescence and live-cell imaging experiments HEK cells were transfected with Lipofectamine 2000 (Invitrogen).

#### Primary cultured rat neurons

Timed pregnant Sprague-Dawley rats were purchased (ENVIGO) and dissected at embryonic day 18. Dissection media (to make 1L) consists of 50 ml 10x HBSS (Gibco), 5 ml Penicillin-Streptomycin (Thermo Fisher Scientific), 5 ml pyruvate (Gibco), 5 ml Hepes (Gibco) 10 mM final, 15 ml of 1M Glucose stock 30 mM final and 420 ml Mill-Q water. Cortical cells were plated onto glass coverslips coated with Poly-L-Lysine hydrobromide (Sigma) at a density of 250K/well (of a 12-well plate) and grown in Neurobasal Plus Medium (Gibco) supplemented with 2% B-27 Plus (Gibco), 2mM Glutamax (Thermo Fisher Scientific), Penicillin-Streptomycin (100U/ml, Thermo Fisher Scientific), and 5% horse serum (Hyclone). Hippocampal cells were plated at a density of 100K/well (of a 12-well plate) in the same manner except with standard Neurobasal Medium and standard B27. For hippocampal cultures, cells were swapped to serum-free media (as above but lacking horse serum) at DIV1 and subsequently fed once a week with serum-free media. For cortical neurons, cells were fed with media containing 1% horse serum and FdU (Sigma) at DIV4, then subsequently fed twice per week with serum free media (Neurobasal Plus Medium and B27 Plus).

#### Primary cultured mouse neurons

P0 Btbd11^F/F^ pups were dissected and plated as above except 5% fetal bovine serum was used in place of horse serum, and that Neurobasal Plus Medium and B27 Plus supplement was used for both hippocampal and cortical cultures. Hippocampal neurons were plated onto Poly-L-Lysine coated coverslips at a density of 200K/well (of a 12-well plate) and cortical neurons were plated at a density of 6.5-7M cells per 10cm plate (also Poly-L-Lysine coated. Cells were fed twice a week with serum free media. AAVs were added to cultures at DIV1 or DIV2. AAV.CMV.PI.EGFP.WPRE.bGH (Serotype 2/9; titer ≥ 1×10^13^ vg/mL), AAV.CMV.HI.eGFP-Cre.WPRE.SV40 (Serotype 2/9; titer ≥ 1×10^13^ vg/mL) and AAV.Syn.NES-jRGECO1a.WPRE.SV40 (Serotype 2/9; titer ≥ 1×10^13^ vg/mL) viruses were purchased from Addgene.

### PSD preparation

Btbd11^F/F^ primary cultured cortical neurons (1×10cm plate per condition) were collected by plate scraping, or one hippocampus (per Gene Trap, vGAT:Btbd11^F/F^ or Btbd11^F/F^ control animal) was dissected, and then homogenized by passage through a 26g needle, 12 times, in homogenization buffer (320mM sucrose, 5mM sodium pyrophosphate, 1mM EDTA, 10mM HEPES pH 7.4, 200nM okadaic acid, protease inhibitor cocktail (Roche)). The homogenate was centrifuged at 800g for 10 minutes at 4°C to yield the P1 and S1 fractions. S1 was further centrifuged at 17,000g for 20 mins at 4°C to yield P2 and S2 fractions. P2 was resuspended in milliQ water, adjusted to 4mM HEPES pH 7.4 from a 1M HEPES stock solution, and incubated with agitation at 4°C for 30 mins. The resuspended P2 was centrifuged at 25,000g for 20 minutes at 4°C to yield LP1 and LS2. LP1 was resuspended in 50mM HEPES pH 7.4, then mixed with an equal volume of 1% triton X-100 and incubated with agitation at 4°C for 15 minutes. The PSD fraction was generated by centrifugation at 32,000xg for 20 minutes at 4°C. The final PSD pellet was resuspended in 1x RIPA buffer. Protein quantification was performed via Bradford assay and samples made up in Laemmli Buffer containing 5% Beta-mercaptoethanol and frozen.

### GST-pulldown experiments

HEK cells (transfected via Ca^2+^ phosphate precipitation) grown to confluence on a 10cm plate were washed 1X with PBS at room temp and then lysed in ice cold lysis buffer (1X Tris Buffered Saline, 1% NP-40, 10mM NaPPi, 10mM NaF, 200nM okadaic acid, 1mM Na_3_VO_4_ and a home-made protease inhibitor cocktail). Cells were scraped from the plate and lysed by rotation for 20 mins at 4°C. Lysates were centrifuged at 17,000g for 15 minutes and the supernatant retained. 20μl of supernatant was kept as 2% INPUT. 30μl of pre-washed Glutathione Sepharose 4B bead slurry (GE Healthcare) was added to each sample and incubated with rotation for 2-hours at 4°C. Beads were washed in spin columns 5X with lysis buffer and then eluted with 30-40ul of Laemmli Buffer containing 5% Beta-mercaptoethanol and frozen or run immediately on western blots.

### Confocal microscopy and image analysis

All confocal imaged was conducted with a Zeiss LSM 880 microscope. Neurons and HEK cells were imaged in 37°C pH 7.4 ACSF (120mM NaCl, 5 mM KCl, 10mM Hepes, 10mM glucose, 2mM CaCl_2_ and 1mM MgCl_2_) in a humidity-controlled chamber. For FRAP experiments (neuron and HEK), the laser power and repetitions needed for successful bleaching was optimized for each experiment, and then kept consistent. Imaged were collected with oil-immersive 40x or 63x objectives, except for the Ca^2+^ imaging experiments in mouse neurons, which was collected using a 10x air objective.

#### Analysis of FRAP data

For FRAP experiments in neurons time-series Z-stacks were acquired with two timepoints acquired before photobleaching. For each dendritic region at least 5 putative synaptic puncta (Psd-95-mCherry puncta) were bleached, ensuring that there were several puncta in the field of view that were not bleached. Images were analyzed in FIJI. Maximum image projections were generated for the timeseries, and a median filter was applied to all channels (1-pixel) and the Azurite cell-fill channel was used to correct for movement during imaging with a rigid-body transformation (MultiStackReg plugin). Regions of interest (ROIs) were drawn around bleached and unbleached puncta as well as region of background (away from the dendritic signal). Background signal was subtracted from the puncta signal, which was then normalized to the average intensity of the unbleached puncta at each timepoint (to account for the low levels of acquisition bleach over time). The signal in the bleached ROIs was then normalized such that the average baseline was centered on 1 and the post-bleach timepoint was 0. For each puncta the estimated maximum recover was estimated with a one phase exponential fit (in GraphPad Prism 9). For comparisons +/-GFP-Btbd11 analysis was performed on decoded data with just the motion-corrected Psd-95-mCherry signal, so the investigator had no knowledge of the experimental condition.

#### Analysis of puncta-to-puncta distance

Imaris 9 (Bitplane) was used to estimate the puncta-to-puncta distance shown in **Figure 1K**. The GFP channel (GFP-Btbd11 knockin) with smoothing (1μm) was used to generate a surface along the dendrite of the knockin cell. This surface was used as a mask for the other channel (*i*.*e*., Psd-95 or Gephyrin) for spot detection and analysis. The spot-detection feature was then used to detect GFP-Btbd11 and Psd-95 or Gephyrin puncta. We then calculated the distance of the nearest Psd-95 or Gephyrin puncta to each GFP-Btbd11 puncta (using the center of the detected spot as the center point of each punctum).

#### Analysis of PV levels with immunofluorescence

Imaris 9 (Bitplane) was used to generate a surface around the cell body of PV-IN using PV immunofluorescence signal with smoothing (2-3μm) and an estimated diameter of 5-10μm. This surface was then used to calculate the average intensity of PV for each cell.

### Cloning and molecular biology

All constructs were generated using HiFi assembly (New England Biolabs). mDlx-Azurite was generated by replacing EGFP with Azurite from the pAAV-mDlx-GFP-Fishell-1 plasmid (Addgene number: 83900). Generation of pORANGE Btbd11 constructs were generated using the pORANGE Cloning template vector (Addgene number: 131471). The guide sequence to target the N-terminus of Btbd11 was: 5-ACGGCGGCTGCAGCATGAAG-3. cDNA for mouse Btbd11 was purchased C-terminal Myc tag (Origene catalog number: MR217199). Using this cDNA as template pCAG-GFP-Btbd11 was generated, removing the C-terminal Myc tag (exposing the PBM of Btbd11). A linker (GGGGSGGGGTR) was added between EGFP and Btbd11. The 5xANK-BTB mutant consisted of the last 512 amino acids of Btbd11 (*i*.*e*., a large N-terminal deletion). pCMV-Psd-95-mCherry point mutants were generated based on the mutations described in (Imamura et al., 2002). pCIS-GST-Btbd11 was generated by subcloning Btbd11 into a pCIS-GST expression vector. The sequence of all constructs was confirmed with DNA sequencing.

### Electron microscopy

Cells grown in 35 mm tissue culture dishes (Falcon 3001) were briefly rinsed with 37 C PBS, then fixed with 2.5 % glutaraldehyde in 100 mM phosphate buffer (Sorenson’s) containing 5 mM MgCl_2_ pH 7.4, for 2.5 hr. at room temperature on a slow rocker. After a 30 min buffer rinse (100 mM phosphate buffer with 3% sucrose and 5mM MgCl_2_), cells were post-fixed in 1% osmium tetroxide in 100 mM phosphate buffer with 5 mM MgCl_2_ at 4°C for 1 hr. in the dark. Samples were then rinsed 100 mM maleate buffer containing 3% sucrose pH 6.2 and en-bloc stained with 2% uranyl acetate (0.22 μm filtered) in the same buffer for 1 hr in the dark. Plates were dehydrated in a graded series of ethanols then infiltrated in Eponate 12 (Pella) overnight without catalyst. The next day cells were further embedded with fresh epon containing 1.5% DMP-30 (catalyst). Culture dishes were cured at 37°C for three days, and further polymerized at 60°C overnight. Cured discs were removed from the plastic dish and 3 mm circles punched out and glued to epon blanks for sectioning. 80 nm thin compression free sections were obtained with a Diatome diamond knife (35 degree). Sections were picked up onto 1×2 mm formvar coated copper slot grids (Polysciences), and further stained with uranyl acetate followed by lead citrate. Grids were examined on a Hitachi H-7600 TEM operating at 80 Kv. Images were digitally captured with an XR-50, 5-megapixel CCD camera (AMT).

### Generation of Btbd11 conditional KO mice

*In vitro* fertilization of C57Blk/6J mice was performed by the Johns Hopkins Transgenic Core using frozen sperm obtained from the European Mutant Mouse Archive (*Btbd11*^*tm1a(EUCOMM)Wtsi*^, https://www.infrafrontier.eu/). Offspring were genotyped as recommended with a common forward primer (5’-3’: TCCTGTCTTAATGCCCCCTG), a wildtype reverse primer (5’-3’: TTCTGGCGGTTCTAAATCCTG) and a mutant reverse primer (5’-3’: TCGTGGTATCGTTATGCGCC). Btbd11 Gene Trap mice were backcrossed with C57Blk/6J animals. To generate Btbd11 conditional knockout mice, Btbd11 Gene Trap mice were crossed with constitutive FLPe-expressing animals. Correct conversion was confirmed with PCR as described by the European Mutant Mouse Archive. Conditional animals were bred to homozygosity. To generate IN specific knockout animals, Btbd11 conditional mice (Btbd11^F/F^) were crossed with vGAT^Cre^ animals (Jackson lab stock: 028862).

### Stereotaxic surgery

#### Virus injection

pAAV-S5E2-dTom-nlsdTom virus (Addgene number: 135630) was packaged by the Janelia Vector core with AAV2/9 serotype (virus titer after 1:1 dilution: 2×10^13^ GC/ml). Stereotaxic surgery was conducted as previously described (Fang et al., 2021). Male and female Btbd11^F/F^ or vGAT:Btbd11^F/F^ mice aged 4-5 weeks were anesthetized with isofluorane, placed into a stereotaxic frame (Kopf) with their body temperature monitored and maintained at 37°C with a closed-loop temperature control system (Kent Instruments). Animals were injected subcutaneously with sterile saline (VetOne; 0.5ml) to maintain hydration and buprenorphine (ZooPharm; 1 mg/kg) and lidocaine (VetOne; 2%) to provide analgesia, with lidocaine injected locally under the skin over the skull. An incision was made to expose the skull with a scalpel, and a craniotomy performed to expose the brain surface (see below for coordinates). Glass pipettes (Drummond Science Company; Wiretrol II) were pulled (Sutter Instruments) and sharpened to a 30° angle (Medical Systems Corp) and used for controlled virus injection with a pneumatic injector (Narishige) at a rate of 100nl/minute. After each injection the pipette was kept in place for 5 minutes before being raised to the next injection depth or being removed slowly from the brain (to prevent backflow of virus). To target the visual cortex the following stereotaxic coordinates and injection volumes were followed, relative to bregma and pia. AP: -3.8, ML: +/-2.6, Z: -450 (200nl) and -300 (200nl). After injection the skin was sutured (Ethicon) and sealed with glue (VetBond). Animals were placed in a heated cage to recover with access to softened food and monitored closely. Animals were left to recover from surgery (and to provide time for virus expression) for at least 12 days before being used for slice electrophysiology experiments.

#### Electrode implant

The initial surgery was performed as described above. Male Btbd11^F/F^ or vGAT:Btbd11^F/F^ mice aged 2-5 months were used for *in vivo* electrophysiology experiments. A craniotomy was made above the V1 in the left hemisphere (AP: level with lambda, ML: 3.2) where a 50μm polyimide-insulated tungsten wire was implanted at a depth of -0.45mm relative to pia. An additional craniotomy was made above the cerebellum just behind lambda and just to the right of the midline for a ground screw. A final craniotomy was made in the right hemisphere (AP: +1.5, ML: 0.5-0.6) for a reference electrode consisting of 125μm stainless-steel coated with PTFE. Wires were connected to a mill-max adaptor with metal pins and the implant secured with light-curable dental cement (3M RelyX). To enable head fixation, a custom-made metal head bar was secured to the skull. Mice were left to recover for 1 week before habituation to handling began. After habituation to handling, mice were habituated to brief periods of head fixation prior to recording sessions. Electrode placements were determined by making electrolytic lesions at the conclusion of experiments which were imaged on DAPI stained brain sections on a fluorescent microscope.

### Slice-physiology

Mice were deeply anaesthetized with isoflurane. Animals irresponsive to toe pinches were decapitated, and brains were rapidly extracted and sectioned using a vibratome (Leica VT-1200). The cutting ACSF contained (in mM) 85 NaCl, 65 sucrose, 25 NaHCO_3_, 10 glucose, 4 MgCl_2_, 2.5 KCl, 1.25 NaH_2_PO_4_, 0.5 CaCl_2_ (pH 7.35, ∼308mOsm). 300 μm horizontal slices containing the prelimbic cortex were collected and recovered for 10min in cutting ACSF at 32°C, after which slices were transferred to ACSF solution containing (in mM) 130 NaCl, 24 NaHCO_3_, 10 glucose, 3.5 KCl, 2.5 CaCl_2_, 1.5 MgSO_4_, 1.25 NaH_2_PO_4_ (pH 7.35, ∼303mOsm). Slices were left to recover for 1h at room temperature before recordings. All ACSF solutions were saturated with 95% O2 and 5% CO2. For recording, a single slice was transferred to a heated chamber (34-35°C) and perfused with ACSF using a peristaltic pump (WPI). tdTomato-expressing cells in the visual cortex were identified on an upright microscope equipped for differential interference contrast (DIC) microscopy (Olympus BX51WI) and LED fluorescence (X-Cite, 120 LED). Whole-cell patch-clamp recordings were made using a MultiClamp 700B amplifier (1kHz low-pass Bessel filter and 10kHz digitization) with pClamp 10.3 software (Molecular Devices). Voltage-clamp recordings were made using borosilicate glass pipets (King Precision Glass Inc., KG-33 ID1.00 OD 1.50) with resistance 2.0-3.0MΩ, filled with internal solution containing (in mM): 117 Cs-methanesulphonate, 20 HEPES, 5 QX-314, 5 TEA-Cl, 4 ATP-Mg, 2.8 NaCl, 1 Na_2_-phosphocreatine, 0.4 EGTA, 0.4 GTP-Na (pH 7.30, ∼290 mOsm). Access resistance was continually monitored throughout recording; cells in which access resistance rose above 20MΩ were excluded from analysis. Membrane potentials were not corrected for junction potentials.

Recordings were conducted in the presence of the GABA receptor antagonist SR-95531 (5 μM) to isolate glutamatergic currents. Electrical stimuli were delivered using a bipolar stimulating electrode (FHC, MX21AES) placed lateral to the recorded cell. Analysis was performed offline using Clampfit (v 10.6) and MiniAnalysis (v 6.0.7, Synaptosoft).

### Ca^2+^ imaging and analysis

Mouse Btbd11^F/F^ hippocampal cultures transduced with AAV-GFP or AAV-GFP-Cre and AAV-jRGECO1a were transferred to ACSF (recipe as above) and allowed to equilibrate for 10-15 mins before being imaged in a temperature and humidity-controlled chamber. Images (512×512 pixels) were acquired with a 10x objective at 4Hz for 60s from multiple regions of interest for each coverslip. In FIJI, each timeseries was projected to a single plane (maximum intensity projection) to aid identifying the soma of neurons. A square/rectangle was drawn at the cell body of neurons and assigned as ROIs. ROIs were then opened on the original timeseries which was median filtered (0.5 pixels). The mean signal intensity was extracted for each neuron in the field of view. Data were then processed with custom written Python scripts using Jupyter notebooks to: 1) convert the signal to represent dF/F assigning F_0_ as the 10^th^ percentile of the signal, 2) calculate the average dF/F of all neurons in each imaging region to capture the synchronous activity, 3) convert the data to a z-score, 4) subtract the minimum value from each timepoint to center the baseline around 0, and 5) use find_peaks and peak_prominences (from scipy library) with parameters [height threshold = 2, distance threshold = 8, prominence threshold = 1] to automatically identify spontaneous activity in the cultures.

### In vivo recordings

Following habituation to handling and head fixation, mice (5 control and 6 knockout) were head restrained and presented with a monitor positioned at 45° with respect to their right eye. As a visual stimulation, mice were presented with a black screen or a gray screen for 90s. Recordings were made with a 32-channel RHD2132 head stage (Intan Technologies, CA, US) via a custom-built adaptor and an Open Ephys acquisition board via an SPI-cable (Intan Technologies). Data were amplified and digitized by the RHD2132 headstage, sampled at 30kHz, and digitally bandpass filtered between 0.1–300 Hz before further processing in Matlab (Mathworks) and custom written scripts in Jupyter notebooks. Data were down sampled to 1kHz and power analyzed with *compute_spectrum* and *specgram* from the Neurodsp and matplotlib libraries, respectively. The mean and peak 55Hz-65Hz power was calculated with presentation of the black or gray screen and converted to a gray/black screen ratio.

### Behavioral testing

Animals were housed in a holding room on a reverse light cycle. Testing was conducted during the dark phase after animals were habituated to handling. Mice were aged between 5 – 6 months at the time of testing. For locomotor activity and spontaneous alternation, we tested 13 control (9 female and 4 male) and 10 knockout (6 female and 4 male) animals. For the MK-801 challenge we used 9 control (5 female 4 male) and 7 knockout (3 female and 4 male) mice.

#### Locomotor activity

Locomotor activity was assessed by placing animals in an open field arena (40×40cm) in the dark and measuring the number of infrared beam breaks during a 90-minute session (San Diego Instruments Inc.). Data were binned into 5-minute periods for analysis.

#### Y-maze spontaneous alternation

Spatial short-term memory was assessed in the Y-maze spontaneous alternation task. Mice were placed at the end of one arm of a Y-maze consisting of three 38cm long arms (San Diego Instruments Inc) and allowed to freely explore the maze for 5 minutes. Animal location was automatically recorded and tracked using Anymaze Tracking Software (Stoelting). The percent of spontaneous alternation was calculated using the equation: % alternation = (total alternations/(total arm entries -2))×100.

#### Locomotor activity with MK-801 challenge

Activity was assessed as above, except the area was illuminated animals received an intraperitoneal injection of saline or MK-801 maleate (0.2mg/kg; Tocris). A within-subject design was followed, and the test was repeated 2-days later with drug assignment reversed. The order of saline vs. MK-801 was counterbalanced for genotype.

### Statistical analysis

GraphPad Prism 9 was used for all statistical analyses. Figures were plotted in Prism 9 or using Python and Jupyter Notebooks. Images were processed in FIJI (Schindelin et al., 2012) and often presented as maximum intensity projections. Images were frequently median filtered to reduce noise. Any adjustment of brightness and contrast was performed uniformly across the image. Figures were assembled in Adobe Illustrator. Details of all statistical tests can be found in **Table S2**. To reduce the chance of bias, FRAP and mEPSC analyses were performed on decoded files and behavioral experiments were conducted by an investigator that was unaware of the genotype of the animals being tested. Data were assumed to follow the normal distribution, but no formal tests of normality were conducted.

